# Fecal metagenomics to identify biomarkers of food intake in healthy adults: Findings from randomized, controlled, nutrition trials

**DOI:** 10.1101/2023.04.10.536271

**Authors:** Leila M. Shinn, Aditya Mansharamani, David J. Baer, Janet A. Novotny, Craig S. Charron, Naiman A. Khan, Ruoqing Zhu, Hannah D. Holscher

## Abstract

**Background:** Undigested components of the human diet affect the composition and function of the microorganisms present in the gastrointestinal tract. Techniques like metagenomic analyses allow researchers to study functional capacity, thus, revealing the potential of using metagenomic data for developing objective biomarkers of food intake.

**Objective:** As a continuation of our previous work using 16S and metabolomic datasets, we aimed to utilize a computationally intensive, multivariate, machine learning approach to identify fecal Kyoto Encyclopedia of Genes and Genomes (KEGG) Orthology (KO) categories as biomarkers that accurately classify food intake.

**Design:** Data were aggregated from five controlled feeding studies that studied the individual impact of almonds, avocados, broccoli, walnuts, barley, and oats on the adult gastrointestinal microbiota. DNA from pre-and post-intervention fecal samples underwent shotgun genomic sequencing. After pre-processing, sequences were aligned and functionally annotated with DIAMOND v2.0.11.149 and MEGAN v6.12.2, respectively. After count normalization, the log of the fold change ratio for resulting KOs between pre-and post-intervention of the treatment group against its corresponding control was utilized to conduct differential abundance analysis. Differentially abundant KOs were used to train machine learning models examining potential biomarkers in both single-food and multi-food models.

**Results:** We identified differentially abundant KOs in the almond (n = 54), broccoli (n = 2,474), and walnut (n = 732) groups (*q* < 0.20), which demonstrated classification accuracies of 80%, 87%, and 86% for the almond, broccoli, and walnut groups, respectively, using a random forest model to classify food intake into each food group’s respective treatment and control arms. The mixed-food random forest achieved 81% accuracy.

**Conclusions:** Our findings reveal promise in utilizing fecal metagenomics to objectively complement self-reported measures of food intake. Future research on various foods and dietary patterns will expand these exploratory analyses for eventual use in feeding study compliance and clinical settings.

## Introduction

The gut microbiome is a complex ecosystem containing over 1,000 bacterial species and their 3.3 million non-redundant genes, which contribute to human health (1,2). While not limited to nutrient metabolism, many of the ways that the intestinal microbiome contributes to host health is through macronutrient metabolism, vitamin production, and bile acid metabolism (3). Metagenomic analyses characterize the microorganisms present in a given sample and their encoded functions, which provide insight into the composition and functional capacity of the microbiome (4). Thus, the use of metagenomics for biomarker discovery is of rising interest (5).

To date, most metagenomic biomarker discovery studies have been specific to disease (6–11). Yet, another promising route for these discoveries is to complement self-reported measures of food intake and compliance with fecal microbial genes and subsequent pathways as objective biomarkers because dietary components affect the fecal microbiome (12). While self-reported food intake and compliance measures are frequently utilized in nutrition studies, they are limited by their reliability and validity (13–17). Therefore, objective biomarkers to complement self-reported measures of food intake, like those identified from metagenomic analyses of fecal samples, are of interest.

The discovery, development, and use of biomarkers of food intake are needed (18–22). Researchers have reported specific microbial genes and pathways associated with food consumption. For example, through daily sampling of the gut microbiome over 17 days, Johnson et al. demonstrated that daily variations in the human gut microbiome relate to food choices (23). In comparing rural and urban Russian gut microbiomes and Japanese and North American gut microbiomes, Tyakht et al. and Hehemann et al., respectively, reported gut microbial signatures attributed to differences in dietary intake (24,25). Furthermore, distinct clusters or “enterotypes” dominated by specific bacteria based on metagenomic sequences have been identified and linked to long-term dietary patterns (26,27). Indeed, the human gut metagenome relates to diet as different dietary components differentially impact gut microbiome composition and function (23–27). Despite the promise of these efforts, more work is needed to fully elucidate the impact of diet on gut microbial composition and function.

Thus, aligned with our previous efforts (28,29), we aimed to develop a proof-of-concept machine learning model to identify microbial metagenomic profiles in fecal samples that could be leveraged as biomarkers of specific food intake. Herein, we describe secondary analyses conducted on data from fecal samples collected at pre-and post-intervention of 5 feeding trials (almonds, avocados, broccoli, walnuts, and whole grains). The purpose of the present investigation was to utilize a computationally intensive, multivariate, machine learning approach to identify fecal Kyoto Encyclopedia of Genes and Genomes (KEGG) Orthology (KO) categories as biomarkers that accurately classify (or predict) participants as either having consumed or not consumed a specific food. Furthermore, the same approach is applied to identify biomarkers that accurately classify the food intake of participants into one of the food groups.

## Subjects and Methods

### Experimental Design

This study utilized data from five separate feeding studies examining almond (30), avocado (31,32), broccoli (33), walnut (34), or whole-grain barley and whole-grain oat (35) consumption in adults (n = 285) between 21 to 75 years of age, which have been briefly summarized in **Table 1**. Briefly, the almond, broccoli, and walnut trials were each complete feeding studies that utilized randomized, controlled, crossover designs. Of note, the original almond trial included five intervention arms: 1) control, 2) whole almonds, 3) whole, roasted almonds, 4) roasted, chopped almonds, and 5) almond butter (30). However, the current effort only included control and chopped almond samples due to cost of analyses and previous efforts revealing statistically significant changes in the gut microbiota composition in chopped versus control samples (36). Further, because the original broccoli trial focused on isothiocyanates metabolism, participants were genotyped for the presence of glutathione S-transferase μ 1 (GSTM1) and glutathione S-transferase θ 1 (GSTT1) genes, which are active in isothiocyanate metabolism (33). Isothiocyanates are derived from broccoli glucosinolates. While glucosinolates have little or no bioactivity, in the presence of myrosinase, they are hydrolyzed to bioactive isothiocyanates and further metabolites. The broccoli used in this study was procured as a single shipment of commercially frozen broccoli, which is blanched and has very little myrosinase activity. Therefore, raw daikon radish was added as a source of plant myrosinase. Charron et al. used this approach because it was impossible to obtain fresh broccoli with consistent glucosinolate content over the duration of the study (33). Thus, while the intervention was technically broccoli and daikon radish, this trial/food will be referred to as “broccoli” throughout the rest of this manuscript for succinctness. The whole grain study was a 6-week, complete feeding, parallel-arm design. The avocado trial was a randomized, parallel-arm, controlled trial that provided one meal daily for 12 weeks. Additional information about the nutrient compositions of the diets for each study are provided in **Supplemental Table 1.** All study procedures were administered in accordance with the Declaration of Helsinki and were approved by the Institutional Review Board of the MedStar Health Research Institute (almond, broccoli, walnut, and whole grains) or the University of Illinois Institutional Review Board (avocado).

**Table 1.** Study design of five studies aggregated for secondary analyses.

| Study | Population |  | Trial Design |  |  |
| --- | --- | --- | --- | --- | --- |
|  | Age, y | BMI, kg/m | Design | Controlled Diet<br>Composition | Intervention Food |
| **Almond**<br>**(n=18)**<br>(30) | 57.0 ± 2.3<br>(25-75 y) | 30.0 ± 1.0<br>(21.9-36.1<br>kg/m<sup>2</sup>) | Five 3-week<br>period<br>randomized,<br>controlled<br>feeding,<br>crossover<br>with 1-week<br>uncontrolled<br>washout<br>periods | Complete feeding (55%<br>carbohydrate, 15% protein,<br>30% fat) | 1.5 servings (42 grams)/day of roasted,<br>chopped almonds (base diet scaled<br>down for isocaloric inclusion) |
| **Avocado**<br>**(n=163)**<br>(32) | 35.0 ± 0.5<br>(25-45 y) | 32.8 ± 0.5<br>(23.9-58.8<br>kg/m<sup>2</sup>) | 12-week<br>investigator-<br>blinded,<br>randomized,<br>controlled,<br>parallel-arm | One daily meal (45%<br>carbohydrate, 15% protein,<br>40% fat) | 175 grams (males) or 140 grams<br>(females) avocado (once daily isocaloric<br>meal) |
| **Broccoli**<br>**(n=18)**<br>(33) | 55.0 ± 1.7<br>(21-70 y) | 28.0 ± 1.2<br>(19.0-36.6<br>kg/m<sup>2</sup>) | Two 18-day<br>period<br>randomized,<br>controlled,<br>crossover<br>with 24-day<br>washout<br>period in<br>adults<br>genotyped for glutathione *S* -transferase $\mu$ 1 (*GSTM1*) and glutathione *S* -transferase $\theta$ 1 (*GSTT1*) gene | Complete feeding (54%<br>carbohydrate, 16% protein,<br>30% fat) | 200 g cooked broccoli with 20 g raw<br>daikon radish/day (added to controlled<br>diet) |
| **Walnut**<br>**(n=18)**<br>(34) | 53.1 ± 2.2<br>(25-75 y) | 28.8 ± 0.9<br>(20.2 -34.9<br>kg/m<sup>2</sup>) | Two 3-week<br>period<br>randomized,<br>controlled<br>feeding,<br>crossover<br>with 1-week<br>washout<br>period | Complete feeding (54%<br>carbohydrate, 17% protein,<br>29% fat) | 1.5 servings (42 grams)/day of walnuts<br>(base diet scaled down for isocaloric<br>inclusion) |
| **Whole grains**<br>**(n=68)**<br>(35) | 52.8 ± 1.3<br>(25-70 y) | 28.2 ± 0.5<br>(18.9-38.3<br>kg/m<sup>2</sup>) | 6-week<br>randomized,<br>double-<br>blinded,<br>controlled<br>feeding,<br>parallel-arm | Complete feeding with 0.7<br>servings (11.2 grams) of<br>whole grains per 1800 kcal<br>(53% carbohydrate, 15%<br>protein, 32% fat) | 4 servings (64 grams) of whole-grain 1)<br>barley or 2) oats per 1800 kcal |

### Sample Collection, DNA Extraction, and Shotgun Genomic Sequencing

All five studies collected fecal samples at the beginning and end of each dietary period. For almond (30), broccoli (33), walnut (34), and whole grains (35), participants were provided with coolers containing dry ice and were instructed to put all fecal samples produced in the coolers immediately after collection. Fecal samples were brought to the center during the participant’s next visit to the center, where they were homogenized and stored at -80°C until they were shipped overnight on dry ice to the University of Illinois for analysis. For avocado (31,32), participants provided fecal samples within 15 min of defecation; upon receipt, samples were homogenized, placed in aliquots, flash frozen, and stored at −80°C.

DNA was isolated from all fecal samples using the PowerSoil DNA Isolation Kit (MoBio Laboratories) according to the manufacturer’s instructions. Stored aliquots of the extracted fecal DNA from the original 16S microbiota work (32,35–38) were utilized for metagenomic sequencing. **Figure 1** outlines the process from DNA extraction through functional annotation. Shotgun genomic DNA libraries were constructed and sequenced at the DNA Services laboratory of the Roy J. Carver Biotechnology Center at the University of Illinois at Urbana-Champaign using the Kapa Hyperprep Sample Preparation Kit (Kapa Biosystems). Briefly, 100 ng high molecular weight DNA was sonicated on a Covaris M220 sonicator to a size of ∼250 bp. After sonication, DNA was blunt-ended, 3′-end A-tailed, and ligated to unique dual-indexed adaptors from Illumina. The adaptor-ligated DNA was amplified by PCR for four cycles with the Kapa HiFi polymerase (Kapa Biosystems). The final libraries were quantitated using Qubit High-Sensitivity DNA (ThermoFisher) and the average size was determined on the Fragment Analyzer (Agilent, CA). The libraries were pooled in equimolar concentration into two pools. Each pool was size selected on a 2% agarose gel for the portion of the DNA library that contained genomic DNA fragments of length 100-350 bp, then evaluated on the Fragment Analyzer. The final pools were diluted to 5 nM concentration and further quantitated by qPCR on a BioRad CFX Connect Real-Time System (Bio-Rad Laboratories). Each pool was loaded on 1 lane of an Illumina NovaSeq 6000 S4 flowcell and sequenced with paired-reads 150nt in length. The FASTQ files were generated and demultiplexed with the bcl2fastq v2.20 Conversion Software (Illumina).

**Figure 1.**
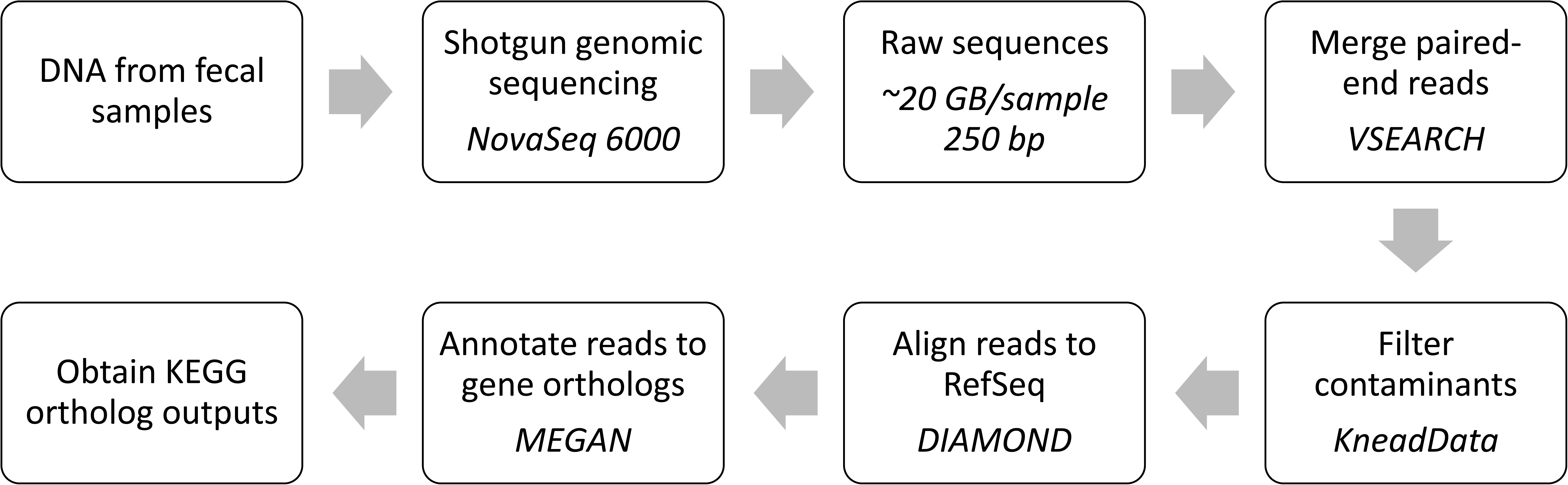
Data workflow from DNA extraction to KEGG functional ortholog counts. An overview of methods from shotgun genomic sequencing, generation of raw sequences, merging paired-end reads, filtering contaminants, aligning reads with DIAMOND, annotating reads in MEGAN, and generation of KEGG orthologs is shown.

### Sequence Pre-processing

All pre-processing steps were performed for each participant’s sequence data, consisting of separate forward and reverse read sequence files for both the pre-intervention and post-intervention timepoints, totaling 4 FASTQ files (samples) per participant.

For each pair of samples, the forward and reverse read FASTQ files were merged into a single-read FASTQ file using the *fastq_mergepairs* function in VSEARCH v2.4.3 (39), a computational tool for pre-processing metagenomic sequences. Each merged FASTQ file was further augmented by concatenating it with the remaining forward reads not merged by VSEARCH. Quality control was then performed on the resulting merged sequence data using KneadData v0.8.0 (40) to separate contaminant host reads from the microbial reads. KneadData removed reads appearing in the *hg37 v0.1* human reference database from each FASTQ file.

### Functional and Taxonomic Annotation

All functional annotation steps were performed for each participant’s merged and cleaned pre-intervention and post-intervention sequence data, totaling 2 FASTQ files (samples) per participant. See **Supplemental Figure 1** for more details on included data.

DIAMOND (double index alignment of next-generation sequencing data) v2.0.11.149 (41) was used in conjunction with the National Center for Biotechnology Information (NCBI) non-redundant (nr) protein reference database (42) to align translated DNA query sequences. The database was downloaded directly from NCBI’s FTP server in June 2021 and formatted using the *makedb* function within DIAMOND. Each sample’s sequences from the merged and cleaned FASTQ file were aligned against the NCBI-nr database, producing a corresponding output DIAMOND alignment archive file. DIAMOND was set to “sensitive” mode, targeting alignments with >40% identity with an e-value of 0.00001.

MEGAN (MEtaGenome ANalyzer) v6.12.2 (43) Ultimate Edition was then used to perform functional analysis of the sequence alignments against the KEGG gene database (44–46). For each sample, the sequence alignments produced by DIAMOND in the previous step were matched to a KEGG ortholog (KO) accession, producing a MEGAN file containing the total count of each KO across each sample. KOs represent common functionalities across orthologous genes in different species based on sequence similarity, enabling the comparison of microbial functional profiles. The MEGAN file was then exported to CSV format for further processing. NCBI taxonomy counts were also exported from MEGAN in a similar fashion.

### Count Normalization

Count normalization was performed on the KO and taxon count table produced by MEGAN6 prior to downstream analysis. First, all counts were normalized using log transformation, offsetting each count by 1 to account for zero-valued data points. To account for intra-study batch effects between participants (e.g., the composition of the microbiome prior to the study), the difference between the pre-and post-intervention log-normalized counts was computed for each sample resulting in two sets of counts for each sample describing the log of the fold change ratio in both KO counts and taxon counts from pre-to post-intervention.

### Differential KEGG Ortholog Abundance and Pathway Enrichment Analysis

Differential KO abundance analysis was conducted individually for each food group by contrasting the normalized KO counts of the food intervention group against its corresponding control group using Student’s t-test (47). Briefly, differential gene expression analysis (DGE or DE for short) is a technique for identifying genes that are statistically different in their expression across the groups being compared (48). KOs were considered differentially abundant if they met a significance threshold of *q < 0.20* after controlling for the false discovery rate (FDR) using the Benjamini-Hochberg procedure (49).

Pathway enrichment analysis was then performed using the *kegga* function in the *limma* R package (50). Pathway enrichment analysis uses the output of a differential expression analysis to describe which broader functional pathways are differentially expressed based on how their constituent genes or KOs are differentially expressed (51). KEGG pathways were tested for over-representation in the set of differentially abundant KOs for each food using Fisher’s exact test (52). Pathways with an uncorrected *P* < 0.05 were considered significantly enriched.

### Machine Learning

We utilized random forests (53) to further examine the relationship between food consumption and changes in functional abundance. For each food group, a *scikit-learn* (54) random forest model with 2000 trees was trained to classify each participant’s study arm (control or treatment) using the normalized KO counts as the covariate. In other words, these KO counts comprise the feature set used in training the model. Only differentially abundant KOs with an absolute mean log-fold change of greater than two were included in the training dataset. Each model was trained and then evaluated in a leave-one-out cross-validated fashion with all model parameters fixed to their default values. The random forest model assigns an importance score (known as the Gini importance score (55)) to each KO describing its relative utility in differentiating the target label i.e., food consumed. KOs with a higher importance are more useful in classifying food intake compared to those with lower importance scores. These importances were extracted from each model to determine the most informative KOs as potential biomarkers in discriminating food consumption.

Finally, we pooled normalized KO counts from each group for classification to examine the impact of food consumption on functional abundance across different food groups. To account for inter-study batch effects (such as varying background diets), we further normalized the data by computing the fold change ratio between each participant’s treatment and control KO counts. Then, we fit a *scikit-learn* random forest model with 2000 trees using the normalized KO counts as the covariate and the food consumed as the outcome. Only KOs considered differentially abundant in at least one of the food groups and with an absolute mean log-fold change of greater than two were included in the training data. Each model was trained and then evaluated in a leave-one-out cross-validated fashion with all model parameters fixed to their default values. KO importance scores were again extracted from each model to determine the most informative KOs in discriminating food consumption.

The pooling of data from various independent studies would typically make the model susceptible to the batch effect, where the training procedure could potentially learn to discriminate the outcome based on variance in the data not influenced by food consumption but rather external factors such as the background diet (28). As each individual in the almond, broccoli, and walnut studies participated in both the food intervention and control arms as part of the crossover design, we utilized a model that accounted for the batch effect (28) by using training data that consisted of the *difference* of the normalized counts between both study arms and endpoints for each individual. This ensured the model was training on data representing the effect of only the food intervention on KO abundance, accounting for the impact of background diets and other external factors. Additionally, it is crucial to understand the interpretations of these findings. While a KO may have a negative log of the fold change ratio, it does not necessarily mean that this KO exhibited lower abundance in absolute terms.

To compare earlier findings on the impact of food consumption on fecal bacteria, we replicated our prior analysis which utilized 16S (V4 region) sequencing to infer microbial abundance with the metagenomic dataset (28). Briefly, the analysis aimed to identify a compact set of microbial biomarkers of specific whole food intake. A marginal screening process was used independently on each food group to select the top 20 most statistically significant microbes when comparing the treatment and control groups within that food’s dataset. This feature set was pooled together and used to train a random forest model to classify which treatment (i.e. almond, avocado, broccoli, walnut, or whole grains) each participant across the treatment groups received. The set of biomarkers was further pared down by pooling the top 10 most important features as ranked by the random forest model for classifying each food group. This compact set of features was then used to train a second random forest to classify which treatment the participants received, demonstrating the effectiveness of the compact set of microbial biomarkers at differentiating whole food intake. Finally, the same random forest model was used to classify participants in the control group to validate that the performance of the model was not heavily influenced by batch effects in which the model is differentiating between differences in background diet or participants in specific studies rather than the effects of the food intake itself. This methodology, originally using taxonomic abundance data derived from 16S sequencing data, was replicated using the NCBI taxonomic data exported from MEGAN. As only a smaller subset of the participants from the original study (n = 285) were included in this analysis, we also replicated the analysis on the original SILVA-annotated (56) 16S data using only this subset of participants.

## Results

The relative abundance of the 20 most variable KOs within each food group before and after both arms of each study are visualized in **Figure 2**. Notably, the almond, broccoli, and walnut groups displayed large shifts in functional composition in the treatment group compared to the control group. In contrast, the avocado and grains groups maintained relatively steady abundances between the treatment and control arms.

**Figure 2.**
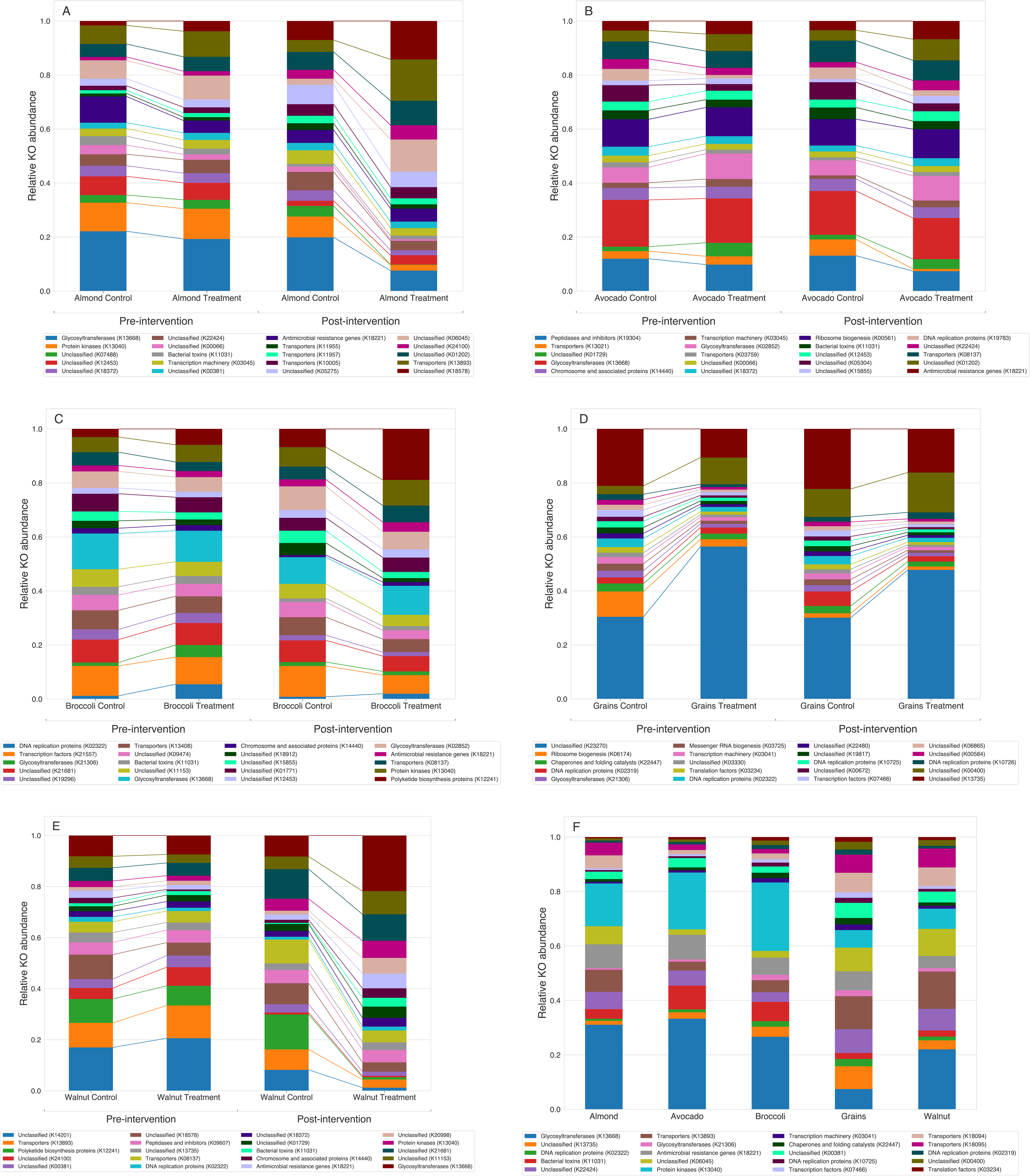
Mean relative abundance of 20 most variable functional orthologs in metabolically healthy adult participants. A) almond, B) avocado, C) broccoli, D) grains, and E) walnut visualize these orthologs before and after undergoing each intervention. Variance was calculated within each food group using KO counts aggregated across all participants in the group and normalized. F) visualizes the most variable orthologs across all food groups prior to each intervention. Mean relative abundance was computed only within a set of 20 most variable orthologs relative to each other. Directional indicator arrows follow changes in mean relative abundance for each ortholog before and after undergoing each intervention type. Bars of the same color represent the same functional ortholog within each of the subpanels A-F. However, the colors do not translate to the same functional ortholog across the different subpanels.

### Differential KEGG Ortholog Abundance and Pathway Enrichment Analysis

The analysis revealed differentially abundant KOs in the almond (n = 54), broccoli (n = 2,474), and walnut (n = 732) groups at a corrected *q-*value of *<* 0.20. No KOs were differentially abundant at this threshold for the avocado, whole-grain barley, or whole-grain oats groups. Therefore, these food groups were excluded from further analysis. **Supplemental Table 2** lists the top 50 most significant differentially abundant KOs found for the almond, broccoli, and walnut groups and their corresponding *q*-values. **Figure 3** shows the number of differentially abundant KOs unique to and shared between each group. Almond and broccoli shared 41 unique differentially abundant KOs, whereas broccoli and walnut shared 551. Almond and walnut shared two unique differentially abundant KOs, manganese-dependent ADP-ribose/CDP-alcohol diphosphatase (K01517) and heparin lyase (K19050). Only two unique KOs, Vitamin B_12_ transporter (K16092) and type III secretion protein R (K03226), were differentially abundant in all three groups.

**Figure 3.**
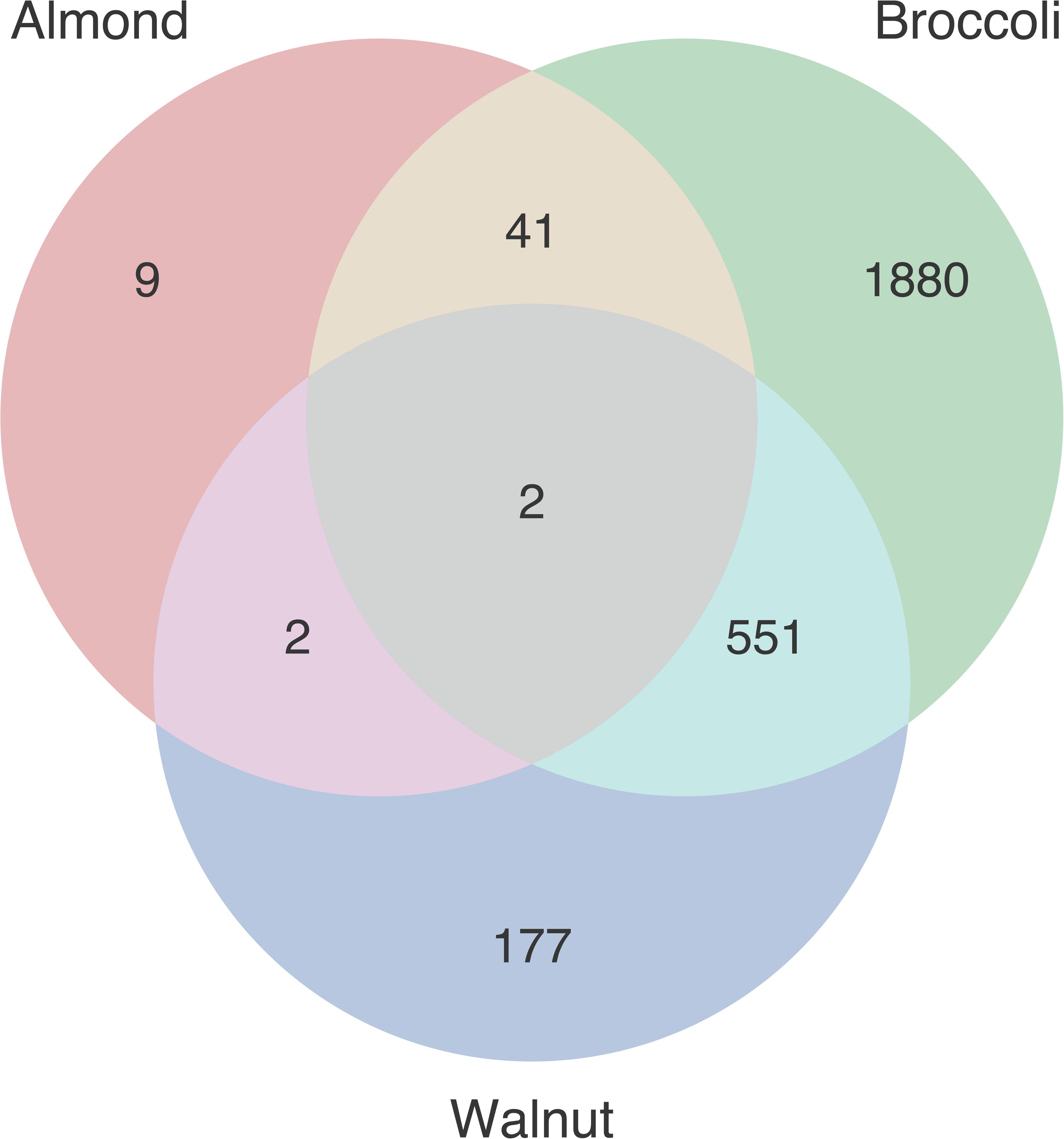
Overlap of differentially abundant KEGG orthologs across almond, broccoli, and walnut. KEGG orthologs (KOs) were considered differentially abundant if they met a significance threshold of q < 0.20. Subset labels indicate the number of KOs differentially abundant in both groups represented by the subset.

Pathway analysis was conducted individually for each food group using over-representation tests by the *kegga* function in the *limma* R package (50). The analysis revealed 4, 59, and 24 pathways with a significant number of constituent differentially abundant KOs in the almond, broccoli, and walnut groups, respectively, at an uncorrected threshold of *P* < 0.05. **Supplemental Tables 3, 4**, and **5** list these pathways, the number of total KOs in each pathway, the number of differentially abundant KOs in each pathway, and the *P* value for the over-representation test.

### Single-food Models

Single-food machine learning models were constructed individually for each food group using the log of the fold change ratio of KO counts for that food group between pre-and post-intervention for both the food intervention and control groups as the covariate and study arm (control or intervention) as the outcome label. The models achieved classification accuracies of 80%, 87%, and 86% for the almond, broccoli, and walnut groups, respectively. The top 10 feature importance scores extracted from each random forest model are shown in **Supplemental Table 6**. The feature importance score distributions for each of the three models all exhibited the “elbow” pattern (57) (**Figure 4A**), i.e., the first few top features show a steep decline in importance, with the subsequent features declining in importance at a slower pace. Of note, the variable importance scores assigned by the random forest model may change slightly each time the model is refit; this numerical instability occurs due to nondeterminism intrinsic to the random forest algorithm.

**Figure 4.**
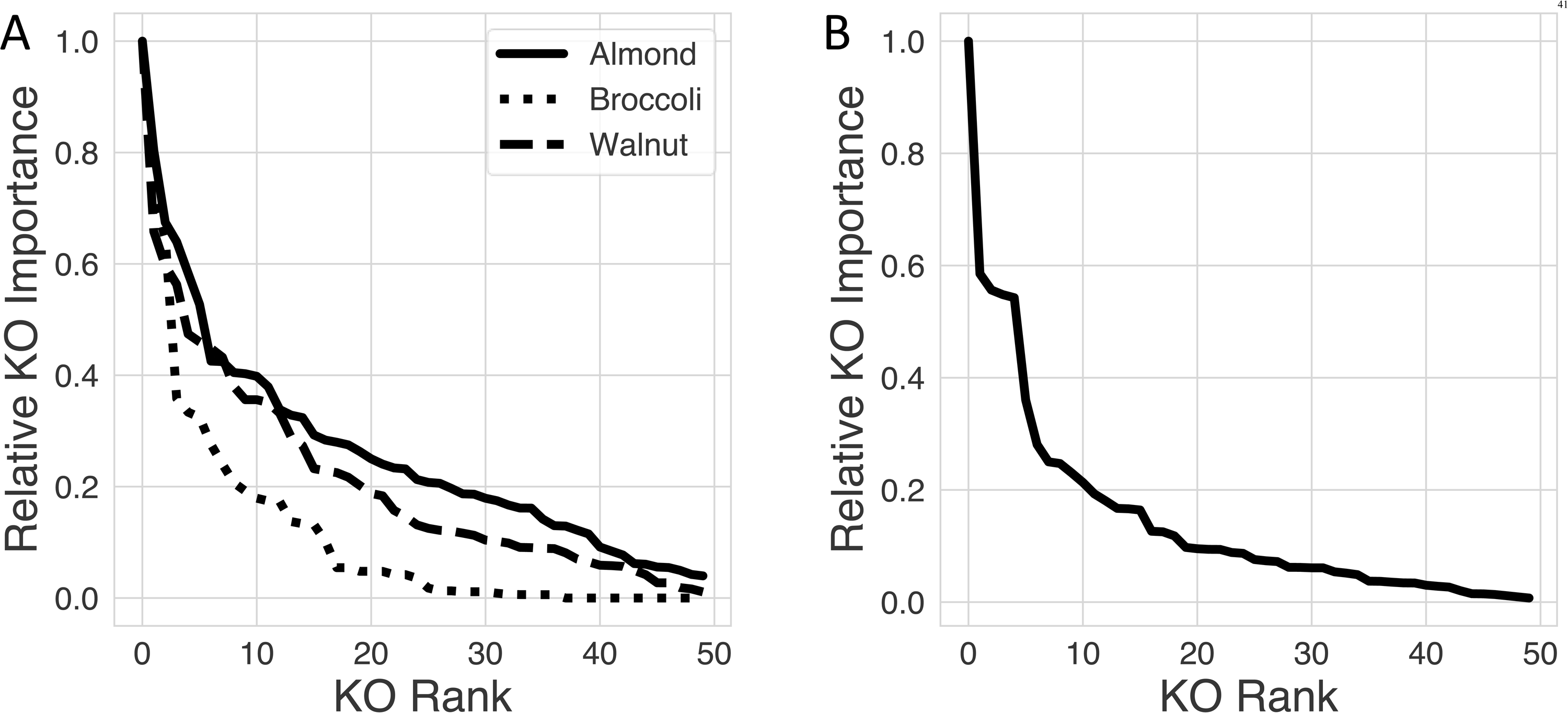
Single-food (A) and multi-food (B) KO importances generated from random forest in almond, broccoli, and walnut. Random forest models were trained to (A) discriminate food intake vs. control for each food group using normalized KO counts and (B) discriminate between almond, broccoli, and walnut intake using normalized KO counts. The top 50 KO importance scores as ranked by the random forest model using the Gini method were extracted and scaled with respect to the most important KO. For almond single-food model at 7 KOs, broccoli single-food model at 3 KOs, and walnut single-food at 9 KOs, the importance scores begin to decline slower than the importance scores before these cutoffs. For the multi-food model, the same trend appears at 10 KOs.

#### Almond

The top 10 KO categories (in rank order) identified by our random forest model for classifying almond treatment versus control consumption resulted in 80% classification accuracy: 1) manganese-transporting P-type ATPase C (K12950), 2) putative colanic acid biosynthesis glycosyltransferase (K13683), 3) a nitrilase involved in tryptophan metabolism (K01501), 4) membrane-bound hydrogenase subunit mbhJ (K18023), 5) RNA polymerase sigma-32 factor (K03089), 6) glycolate oxidase iron-sulfur subunit (K11473), 7) probable lipoprotein NlpC (K13695), 8) serine/threonine-protein kinase PpkA (K11912), 9) RHH-type transcriptional regulator, proline utilization regulon repressor/proline dehydrogenase/delta 1-pyrroline-5-carboxylate dehydrogenase (K13821), and 10) an outer membrane lipoprotein carrier protein (K03634).

#### Walnut

For walnut’s 86% classification accuracy, the top 10 KO categories included four ATP-binding cassette (ABC) transporters (1) K10562, 2) K16013, 3) K10559, and 9) K05658), 4) spore coat-associated protein N (K06336), 5) an uncharacterized protein (K09145), 6) L(+)- tartrate dehydratase alpha subunit (K03779), 7) thiol-activated cytolysin (K11031), 8) heparin lyase (K19050), and 10) an RCC1 and BTB domain-containing protein (K11494).

#### Broccoli

Finally, the ten broccoli KO categories (in rank order) included 1) heptosyltransferase II (K02843), 2) probable lipoprotein NlpC (K13695), 3) 4-phytase/acid phosphatase (K01093), 4) MFS transporter/FSR family fosmidomycin resistance protein (K08223), 5) β-barrel assembly-enhancing protease (K01423), 6) 3-deoxy-D-manno-octulosonate 8-phosphate phosphatase (K03270), 7) MFS transporter/NHS family xanthosine permease (K11537), 8) D-glycero-beta-D-manno-heptose-7-phosphate kinase (K21344), 9) tyrosine-protein kinase Etk/Wzc (K16692), and 10) phosphomannomutase/phosphoglucomutase (K15778), resulting in 87% classification accuracy.

### Mixed-food model

The mixed-food machine learning model was constructed using the difference of the log of the fold change ratio of KO counts for each food group between pre-treatment and post-intervention between the control and treatment arms of each food group and treatment arm (almond, broccoli, or walnut) as the outcome label. The overall mixed-food random forest achieved a classification accuracy of 81%. The feature importance distribution of the mixed-food model was similar to that of the single-food models, exhibiting an “elbow”-shaped curve where the first few features saw a steep drop-off in importance while the following features experienced a less steep decline (**Figure 4B**). The top 25 feature importances were extracted from the mixed-food model (**Supplemental Table 7**) to encapsulate the most useful KOs while keeping the set compact and retaining ease of visualization.

The difference of the log of the fold change ratio of the top 25 most important features extracted from the model was visualized across the three groups (almond, broccoli, and walnut) in a heatmap (**Figure 5**), demonstrating that these features show potential as biomarkers of dietary intake as differences were seen when comparing treatment to respective control groups within each food. As shown in Figure 5, 15 KOs were increased in walnut treatment compared to control. In contrast, the 15 KOs enriched in the walnut treatment compared to control decreased in treatment compared to control for almond samples. Of the 15 KOs increased in walnut treatment compared to control, 12 decreased in broccoli treatment compared to control, whereas three KOs also increased in broccoli treatment compared to control. On the other hand, five KOs increased in almond and broccoli treatment compared to control but decreased in walnut treatment compared to control. Finally, five KOs increased in broccoli treatment compared to control, but decreased in almond and walnut treatment compared to their respective controls.

**Figure 5.**
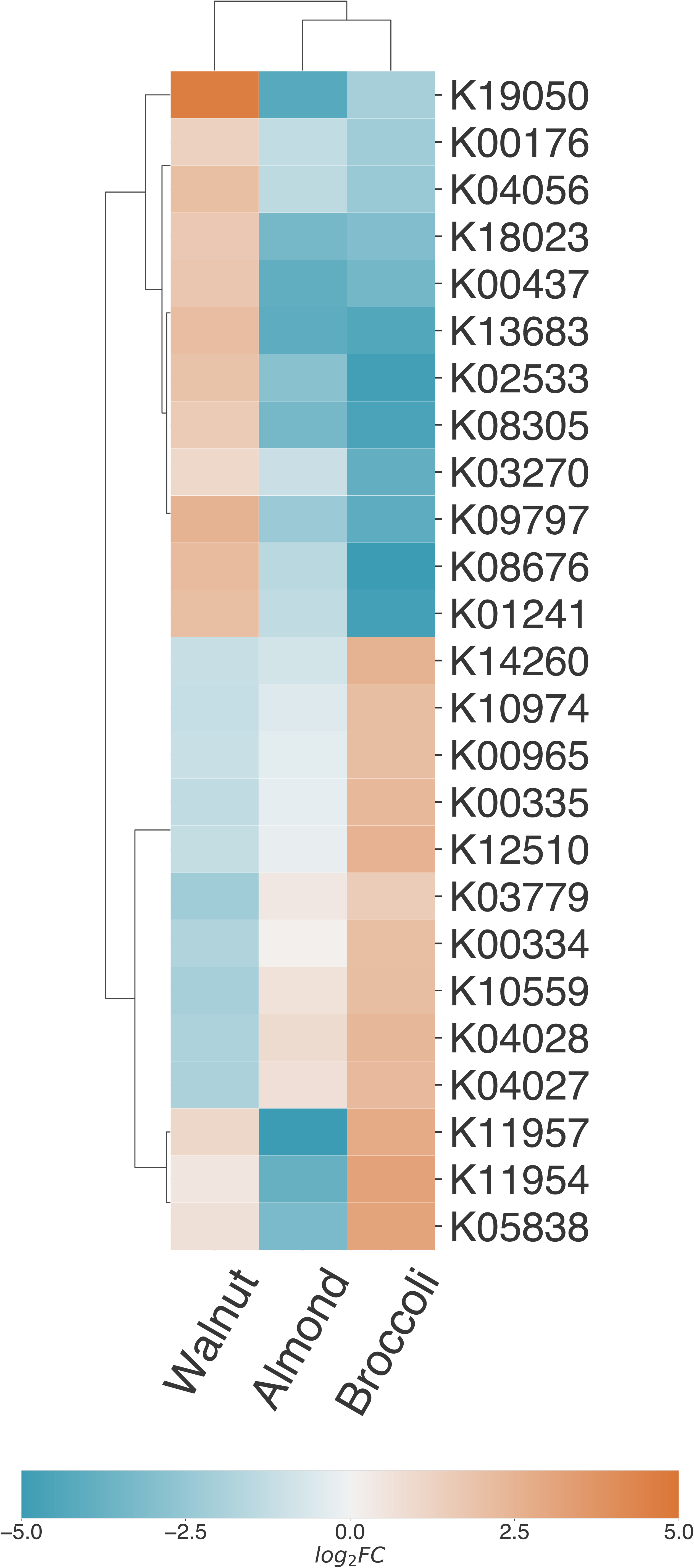
Heat map of almond, broccoli, and walnut top 25 features selected by multi-food random forest model. A random forest model was trained to discriminate between almond, broccoli, and walnut intake using normalized KO counts. The top 25 most important KOs as ranked by the random forest model using the Gini method were extracted. Orange boxes indicate an increased mean fold change from pre-to post-intervention in each food group’s treatment group with respect to control, whereas blue boxes indicate a decreased fold change. The darker the color, the higher the magnitude of change for that KO. The dendrogram (black bars) was generated using Euclidean distance metric for both study groups and the individual KOs. Bars across the top and y-axis show how variables cluster together. Items that are in the same cluster are more similar (i.e., across the top, hierarchical clusters show which foods have similar patterns of fold change across the KO category, and across the y-axis the clusters show which KOs have similar patterns of fold change across the food groups). This clustering visualizes why a given KO was ranked as important by the model – KOs with a different shift in expression within a specific food group only serve as a useful biomarker for consumption of that food group.

### Replication of previous 16S methods on NCBI-nr data

Normalized taxonomic counts relying on annotations of whole genome shotgun sequences from the NCBI-nr database in the current effort were used in place of annotations of 16S sequences from the SILVA database to replicate our previous work (28). As detailed in the methods, features (i.e., 86 microbes) were selected by pooling the top 20 most important features from each food group using the Kruskal-Wallis test (58). These features were used to train a random forest model to classify which food intervention each participant received. The features (i.e. microbial biomarkers) and their importances across all the foods are listed in **Table 2**. The top 10 most important features for each food group as ranked by the initial random forest model were extracted, resulting in 29 unique features. This final compact dataset was then used to train a second random forest model to classify food intake. This model achieved per-class balanced classification accuracies of 69%, 80%, 87%, 83%, and 92% on the almond, avocado, broccoli, walnut, and whole grains groups respectively. The overall accuracy was 74% (AUC = 0.93) above the no information rate of 29% with *P* < 0.05. The confusion matrix for the model is shown in **Supplemental Table 8**. When the same model was used to classify participants in the control arm of each study, it achieved per-class balanced accuracies of 44%, 71%, 48%, 79%, and 66% on the almond, avocado, broccoli, walnut, and whole grains groups, respectively, for an overall balanced accuracy of 41% above the no information rate of 24% with *P* < 0.05 and an AUC of 0.69. The confusion matrix for this model is also shown alongside the prior results in Supplemental Table 8. Finally, when retraining and classifying on the original SILVA-annotated 16S dataset using the subset of participants also included in this study, the model achieved per-class balanced classification accuracies of 61%, 73%, 77%, 91%, and 83% on the almond, avocado, broccoli, walnut, and whole grains groups, respectively, for an overall balanced accuracy of 67% above the no information rate of 29% with *P* < 0.05 and an AUC of 0.89. The microbial biomarkers with their feature importances extracted from this model are listed in **Supplemental Table 9**.

**Table 2.**
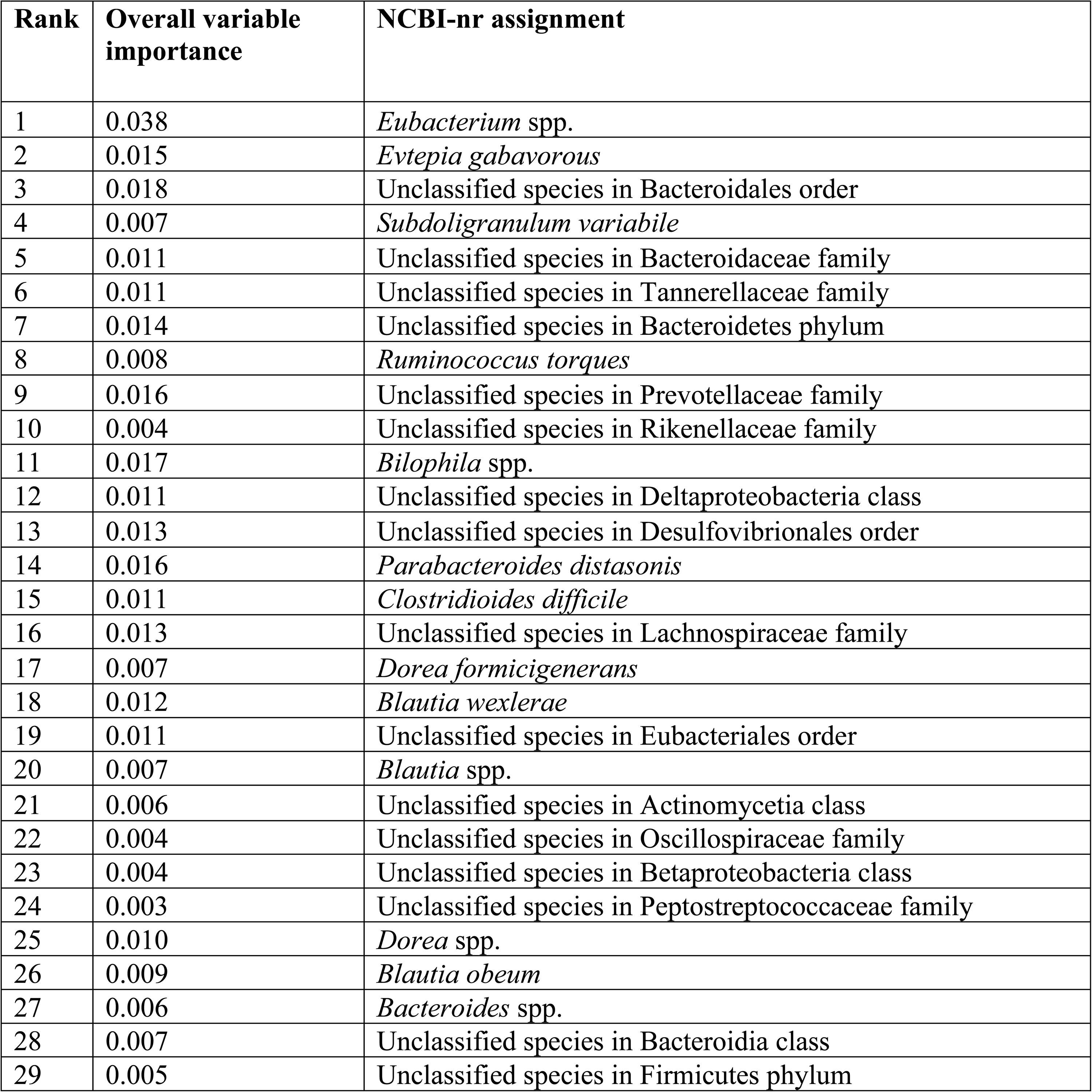
Microbial biomarkers using the top fecal metagenomic NCBI-nr annotated species from metabolically healthy adults who consumed 5 foods.

Across both the NCBI-nr and SILVA 16S models, classification of the control groups using the final random forest model trained on the treatment groups resulted in poor classification accuracy (Supplemental Table 8). The NCBI-nr trained model achieved an overall accuracy of 41% (AUC = 0.68) and the SILVA 16S model achieved an overall accuracy of 40% (AUC = 0.67). As both models performed poorly, we have more confidence that their accuracies on the treatment groups were not entirely due to overfitting or batch effects (28), such as the study site or background diet, which differed across the food groups. However, both models still performed better than the no information rates with *P* < 0.05, indicating the presence of at least some batch effects.

## Discussion

Herein, we report fecal microbial KO categories and subsequent metabolic pathways associated with individual food intake (i.e., almond, broccoli, and walnut). This effort, which utilized random forest models to identify food intake biomarkers, revealed high classification accuracy of almond, broccoli, and walnut intake, both individually (compared to respective controls) and in a mixed-food model (almond versus broccoli versus walnut). Further, we identified differentially abundant KOs across all three food groups. Of note, the most promising findings were produced from data in our three randomized, controlled, crossover, complete-feeding trials compared to our two parallel-arm trials. These findings reveal the promise of metagenomic data from rigorously designed research efforts in establishing fecal KOs as biomarkers of food intake to objectively complement self-reported food measures and study compliance. Of note, differentially abundant KOs were considered statistically significant at a conservative *q*-value of 0.20 due to the exploratory nature of our analyses. For pathway analysis, *kegga* intentionally provides uncorrected *P* values as KOs can be part of multiple pathways, and thus, FDR correction would be overly conservative (49). However, we chose a more conservative *P* = 0.05 to account for the lack of FDR correction.

With approximately 40 grams of dietary carbohydrates, 12-18 grams of protein, and 5% of dietary lipids bypassing human digestion each day (12), these food components are available for digestion by over 1,000 bacterial species and their 3.3 million non-redundant genes (2), some of which encode approximately 16,000 carbohydrate-active enzymes (CAZymes) (59). Because of this enzymatic capacity, there is interest in the relationship between dietary intake and fecal microbial genes (23–27). Johnson et al. demonstrated that the human gut microbiome relates to food choices rather than individual nutrients, noting that while food-microbe interactions are highly personalized, it is likely that specific dietary compounds will have consistent effects on certain bacterial strains and metabolic pathways (23). In considering geographical differences, Tyakht et al. reported that Russians in several rural regions had gut microbiomes dominated by bacteria from the Firmicutes and Actinobacteria phyla, including *Ruminococcus bromii* and *Eubacterium rectale*, which can metabolize resistant starch (24). Therefore, it was hypothesized that these microbiome signatures were related to the consumption of conventional staple foods in rural Russia, including starch-rich bread and potatoes. The lack of these starch-metabolizing microbes in Western cohorts was likely due to reduced consumption of resistant starch in these regions. Thus, Tyakht et al. attributed differences in the rural versus urban gut microbiomes to multiple factors, including diet. In the Japanese population, Hehemann et al. demonstrated that genes encoding porphyranases, CAZymes involved in the degradation of red algae, are present in the Japanese gut microbiome, but absent in that of North Americans. As seaweeds are an important component of the Japanese diet, the Japanese human gut microbiome’s acquisition of these CAZymes stands to reason (25). Arumugam et al. identified three distinct clusters or “enterotypes” based on metagenomic sequences that were dominated by Bacteroides, Prevotella, or Ruminococcus with enrichment for specific gene functions (26). Wu et al. confirmed that long-term dietary patterns were the primary predictor of an individual’s enterotype. Further, the Bacteroides enterotype was associated with a Western diet, high in proteins and fat, while the Prevotella enterotype was associated with consumption of plant fiber (27). Finally, Turnbaugh et al. demonstrated that changing from a low-fat, plant polysaccharide-rich diet to a high-fat/high-sugar “Western” diet changed microbiome gene expression in humanized gnotobiotic mice, further supporting the adaptability of the composition of the human gut microbiome, and, therefore, function in relation to diet (60).

In the present study, manganese-dependent ADP-ribose/CDP-alcohol diphosphatase (K01517) and heparin lyase (K19050) were differentially abundant in both almond and walnut treatment samples, which may play roles in immune cell signaling and phospholipid biosynthesis (61) and the cleavage of glycosidic bonds in polysaccharides (62), respectively. Further, vitamin B_12_ transporter, BtuB (K16092), and type III secretion protein R (K03226), were differentially abundant in all three groups. Like humans, gram-negative bacteria require essential nutrients and thus have mechanisms to obtain cofactors, such as cobalamin, from external sources (63). While many researchers focus on type III secretion systems to discover antimicrobial therapies against pathogens, these systems are also present in symbiotic bacteria as they are highly conserved across bacterial species, playing an important role in various cellular activities by delivering effector proteins to targeted eukaryotic cells (64). Of note, butanol dehydrogenase (K00200), an enzyme involved in microbial fermentation, was differentially abundant in fecal bacteria of participants in the walnut intervention (65). The breadth of functional potential demonstrated in these four differentially abundant KOs alone highlights the wide variety of roles that the gut microbiome plays throughout the body, reflecting the potential for targeting compositional and, therefore, functional changes through diet.

Examining the top 10 KO categories identified by our random forest model for almond, broccoli, and walnut individually, we see features related to genetic information processing, signaling and cellular processes, and carbohydrate, amino acid, and vitamin pathways. Of note, broccoli achieved the highest classification accuracy of the three foods examined here compared to our previous work utilizing 16S rRNA bacterial sequence data, in which broccoli was our lowest performing category (28). Similar to our single-food model, the majority of features identified as important by our random forest model in our multi-food analyses are protein families related to various metabolic processes, genetic information processing, and signaling and cellular processes, supporting evidence that diet may alter the activity and function of the human intestinal microbiome (60,66). These findings highlight the importance of using multiple-omics techniques in biomarker discovery.

Random forest models are well-suited for metagenomics classification and biomarker selection tasks (67). Yatsunenko et al. compared microbiome functional profiles across demographics using random forest models trained on KEGG enzyme data to discriminate between age groups and geographical locations (68). Random forests easily generalize from binary problems to multi-class problems, unlike some other types of supervised models, such as logistic regression and support vector machines (SVM). Additionally, random forests have a lower tendency to overfit when compared to SVM models and are uniquely effective in classifying datasets with smaller sample sizes (69,70). Finally, random forests can intrinsically inform biomarker discovery by assigning importance scores to input features without relying on external feature selection tools. As a high score indicates the KO was useful in classifying the food, KOs with high feature importance scores could be promising biomarker candidates.

When comparing the current effort’s NCBI-nr taxonomic annotations with the previous effort’s SILVA annotations of 16S sequences (28), the classification accuracies across the food groups were similar among datasets, with the exception of the balanced accuracy for classifying broccoli increasing from 11% with 16S analysis to 87% with metagenomic analysis (28). However, these results are difficult to directly compare to the previous 16S work (28) due to the differing sample sizes (340 data points (difference in pre-and post-intervention) in the previous 16S work; 187 data points in current report). To more directly compare the metagenomic and 16S work, we examined the results after replicating the analysis on the subset of 187 samples present in both datasets (Supplemental Table 8). When comparing the two analyses performed on the same samples, the model trained on the NCBI-nr metagenomic taxonomic data demonstrated higher classification accuracy of food intake across all food groups than the model trained on the SILVA 16S taxonomic data. When comparing the subsetted 16S model with 187 data points to the original with 340, classification of the almond group performed poorly compared to our original 16S efforts (62% vs 76% accuracy) (28), demonstrating the negative effect of the reduced sample size on this analysis. Furthermore, classification of the broccoli group using 16S taxonomic data on the subset of samples exhibited higher classification accuracy compared to our 16S original efforts (77% vs 11%) (28), indicating that the increased accuracy of the current effort may be inflated due to overfitting or elimination of certain samples. Because of the potential for more accurate taxonomic profiling resulting from the higher resolution of gene sequencing in metagenomics analysis compared to 16S and thus, more insightful biomarkers, researchers may choose to utilize metagenomics data when possible.

From a microbial biomarkers standpoint, in comparing the current effort’s NCBI-nr taxonomic annotations with the previous effort’s SILVA annotations of 16S sequences (28), *Parabacteroides distasonis* and species within the *Lachnospiraceae*, *Subdoligranulum*, and *Bacteroides* genera appeared in both our current (Table 2) and previous efforts (28). Further, species within the *Dorea* and *Ruminococcus* genera, which appeared as important in original efforts (32,38), were identified by the current random forest model. Unique to the current effort, the butyrate-producing genus, *Eubacterium* (71), was deemed important by our random forest model. *Eubacterium* spp. are also involved in bile acid and cholesterol metabolism (72). Finally, *Blautia wexlerae* and *Blautia obeum* were also identified as important features by our random forest model. *Blautia* plays is involved in various metabolic diseases, inflammatory diseases, and biotransformation with recent interest in its potential probiotic properties (73). The consistencies between our previous 16S (28) and current metagenomic effort reveal promise in our ability to identify fecal microbes as objective biomarkers of food intake. However, there are differences in some of our current findings when comparing to our previous effort (28). For example, *Roseburia* was enriched with almond (36) and walnut (38) consumption, and multiple *Roseburia* species were identified as a potential bacterial (16S) biomarkers (28). However, *Roseburia* was not selected by our current metagenomic taxa model. This discrepancy points to the limitations in the microbiota molecular methods, namely primer bias (74). Thus, when feasible, shotgun genomic sequencing should be utilized (75).

Our works reveals promise in the utility of metagenomics data as food biomarkers, which underscores the importance of including metagenomic endpoints as primary outcomes in future studies. While the present study is strengthened by the use of state-of-the-art bioinformatics techniques and fecal samples from three randomized, controlled, crossover, complete-feeding trials, our two parallel-arm trials did not perform well. Specifically, the avocado and whole grain studies utilized parallel-arm designs, which likely contributed to the lack of signal in our differential KO abundance and pathway enrichment analysis as participants were not able to serve as their own control as in the cross-over trials. The intervention approach (i.e., single meal compared with all other trials being complete feeding) is another important consideration—the lack of findings in our avocado data was likely also affected by the feeding intervention being less controlled. With appropriate primary endpoints and large enough sample sizes, parallel-arm studies could be used to identify fecal microbiome biomarkers. In fact, parallel-arm studies can prevent carryover effects between conditions that can affect causal inferences (76,77).

Although metagenomic data provides insight into functional capacity, metabolomics and transcriptomic studies are needed to assess metabolic activity and active genes. Thus, future work should utilize multi-omics analyses, when possible. As technologies in the field advance, it is likely that the cost of metagenomic analyses will decrease in a similar manner to that of 16S sequencing, making metagenomic and multi-omics analysis more feasible. The National Institute of Health’s Nutrition for Precision Health, powered by the All of Us Research Program *All of Us Research Program*, which will incorporate fecal metagenomic and metabolomic analyses, is one example of initiatives that are likely to help spur molecular and computational advancements that may ultimately drive down costs while also increasing the practicality of these types of analyses.

While the gut microbiome expresses many functional genes involved in core metabolic pathways across healthy individuals (26,78), inter-individual variability must also be considered. Future randomized, controlled, crossover, complete feeding trials should also include dose-response of specific foods within a greater number of individuals. Specific to broccoli, future work is needed to examine multi-omic outcomes in raw and cooked broccoli consumption alone, as the current effort provided cooked broccoli in combination with daikon radish. Also, host genetic factors (e.g., SNPs related to absorption and metabolism of dietary components) should be integrated into future trials to better understand host-microbe interactions and improve algorithms used to predict responses to foods and dietary patterns (79). As researchers continue to elucidate the relationship between diet and the gut microbiome and identify microbial genes and pathways as biomarkers of food intake, these outcomes can be examined in observational trials and eventually used in clinical and research settings as compliance measures to complement self-reported measures of intake and advance the field of personalized nutrition.

In summary, using metagenomics data and machine learning, we reveal promise in the feasibility of fecal KO categories as objective biomarkers of food intake. These findings provide groundwork for uncovering additional objective biomarkers of food intake. With future work and integration of -omics data, biomarkers like the ones identified from this effort can be applied in feeding study compliance and clinical settings.

## Abbreviations

CAZymes: carbohydrate-active enzymes
DIAMOND: Double Index AlignMent Of Next-generation sequencing Data
FDR: false discovery rate
GSTM1: glutathione S-transferase μ 1
GSTT1: glutathione S-transferase θ 1
KEGG: Kyoto Encyclopedia of Genes and Genomes
KO: Kyoto Encyclopedia of Genes and Genomes Orthology
MEGAN: MEtaGenome ANalyzer
NCBI: National Center for Biotechnology Information
nr: non-redundant
rRNA: ribosomal RNA
SVM: support vector machine.

## Data availability

All of the code used for the analyses and a mock dataset can be accessed from Github at https://github.com/holscher-nhml/usda-path-metagenomics.

## Acknowledgements

We thank Alvaro G. Hernandez and Chris L. Wright of the DNA Services laboratory of the Roy J. Carver Biotechnology Center at the University of Illinois at Urbana-Champaign for constructing and sequencing DNA libraries for our metagenomic analyses. Further, we thank Melisa Bailey, Heather Guetterman, Jennifer (Kaczmarek) Burton, Annemarie Mysonhimer, Andrew Taylor, and Sharon Thompson for their technical assistance with processing the fecal samples in the primary studies. The authors’ responsibilities were as follows—DJB, JAN, CSC, NAK, HDH, and RZ: designed the research; LMS: conducted the research; AM: analyzed the data and performed the statistical analysis; LMS and AM: wrote the first draft of the paper; HDH and RZ: critically reviewed and edited the manuscript and had primary responsibility for the final content; and all authors: read and approved the final manuscript.

## References

1. Ursell LK, Metcalf JL, Parfrey LW, Knight R. Defining the human microbiome. Nutr Rev 2012;70:S38.

2. Qin J, Li R, Raes J, Arumugam M, Burgdorf KS, Manichanh C, Nielsen T, Pons N, Levenez F, Yamada T, et al. A human gut microbial gene catalogue established by metagenomic sequencing. Nature 2010;464:59–65.

3. Yadav M, Verma MK, Chauhan NS. A review of metabolic potential of human gut microbiome in human nutrition. Arch Microbiol 2018;200:203–17.

4. Claesson MJ, Clooney AG, O’Toole PW. A clinician’s guide to microbiome analysis. Nat Rev Gastroenterol Hepatol 2017;14:585–95.

5. Segata N, Izard J, Waldron L, Gevers D, Miropolsky L, Garrett WS, Huttenhower C. Metagenomic biomarker discovery and explanation. Genome Biol England; 2011;12:R60.

6. Coker OO, Liu C, Wu WKK, Wong SH, Jia W, Sung JJY, Yu J. Altered gut metabolites and microbiota interactions are implicated in colorectal carcinogenesis and can be non-invasive diagnostic biomarkers. Microbiome 2022;10:1–12.

7. Manichanh C, Rigottier-Gois L, Bonnaud E, Gloux K, Pelletier E, Frangeul L, Nalin R, Jarrin C, Chardon P, Marteau P, et al. Reduced diversity of faecal microbiota in Crohn’s disease revealed by a metagenomic approach. Gut 2006;55:205–11.

8. Laske C, Müller S, Preische O, Ruschil V, Munk MHJ, Honold I, Peter S, Schoppmeier U, Willmann M. Signature of Alzheimer’s Disease in intestinal microbiome: Results from the AlzBiom study. Front Neurosci 2022;16.

9. Qin J, Li Y, Cai Z, Li S, Zhu J, Zhang F, Liang S, Zhang W, Hansen T, Sanchez G, et al. A metagenome-wide association study of gut microbiota in type 2 diabetes. Nature 2012;490:55–60.

10. Karlsson FH, Fåk F, Nookaew I, Tremaroli V, Fagerberg B, Petranovic D, Bäckhed F, Nielsen J. Symptomatic atherosclerosis is associated with an altered gut metagenome. Nat Commun 2012;3.

11. Nagata N, Nishijima S, Kojima Y, Hisada Y, Imbe K, Miyoshi-Akiyama T, Suda W, Kimura M, Aoki R, Sekine K, et al. Metagenomic identification of microbial signatures predicting pancreatic cancer from a multinational study. Gastroenterology 2022;163.

12. Duncan SH, Flint HJ, Sheridan PO, Scott KP, Gratz SW. The influence of diet on the gut microbiota. Pharmacol Res 2012;69:52–60.

13. Schatzkin A, Subar AF, Moore S, Park Y, Potischman N, Thompson FE, Leitzmann M, Hollenbeck A, Morrissey KG, Kipnis V. Observational epidemiologic studies of nutrition and cancer: the next generation (with better observation). Cancer Epidemiol Biomarkers Prev 2009;18:1026–32.

14. Freedman L, Potischman N, Kipnis V, Midthune D, Schatzkin A, Thompson F, Troiano R, Prentice R, Patterson R, Carroll R, et al. A comparison of two dietary instruments for evaluating the fat-breast cancer relationship. Int J Epidemiol 2006;35:1011–21.

15. Rennie K, Coward A, Jebb S. Estimating under-reporting of energy intake in dietary surveys using an individualised method. Br J Nutr 2007;97:1169–76.

16. Poslusna K, Ruprich J, de Vries J, Jakubikova M, van’t Veer P. Misreporting of energy and micronutrient intake estimated by food records and 24 hour recalls, control and adjustment methods in practice. Br J Nutr 2009;101 Suppl.

17. Kipnis V, Midthune D, Freedman L, Bingham S, Day N, Riboli E, Ferrari P, Carroll R. Bias in dietary-report instruments and its implications for nutritional epidemiology. Public Health Nutr 2002;5:915–23.

18. Meyers LD, Suitor CW. Dietary reference intakes research synthesis: workshop summary. National Academies Press 2007.

19. Raiten DJ, Namasté S, Brabin B, Combs GJ, L’Abbe MR, Wasantwisut E, Darnton-Hill I. Executive summary--Biomarkers of nutrition for development: Building a consensus. Am J Clin Nutr 2011;94:633S–50S.

20. Maruvada P, Lampe JW, Wishart DS, Barupal D, Chester DN, Dodd D, Djoumbou-Feunang Y, Dorrestein PC, Dragsted LO, Draper J, et al. Perspective: Dietary biomarkers of intake and exposure-exploration with omics approaches. Adv Nutr 2019;11:200–15.

21. Nogal B, Blumberg JB, Blander G, Jorge M. Gut microbiota–informed precision nutrition in the generally healthy individual: are we there yet? Curr Dev Nutr 2021;5.

22. Mandal R, Cano R, Davis CD, Hayashi D, Jackson SA, Jones CM, Lampe JW, Latulippe ME, Lin NJ, Lippa KA, et al. Workshop report: Toward the development of a human whole stool reference material for metabolomic and metagenomic gut microbiome measurements. Metabolomics 2020;16:119.

23. Johnson AJ, Vangay P, Al-Ghalith GA, Hillmann BM, Ward TL, Shields-Cutler RR, Kim AD, Shmagel AK, Syed AN, Walter J, et al. Daily sampling reveals personalized diet-microbiome associations in humans. Cell Host Microbe 2019;25:789–802.e5.

24. Tyakht A v, Kostryukova ES, Popenko AS, Belenikin MS, Pavlenko AV, Larin AK, Karpova IY, Selezneva O v, Semashko TA, Ospanova EA, et al. Human gut microbiota community structures in urban and rural populations in Russia. Nat Commun 2013;4:2469.

25. Hehemann JH, Correc G, Barbeyron T, Helbert W, Czjzek M, Michel G. Transfer of carbohydrate-active enzymes from marine bacteria to Japanese gut microbiota. Nature 2010;464:908–12.

26. Arumugam M, Raes J, Pelletier E, Paslier D le, Yamada T, Mende DR, Fernandes GR, Tap J, Bruls T, Batto JM, et al. Enterotypes of the human gut microbiome. Nature 2011;473:174–80.

27. Wu GD, Chen J, Hoffmann C, Bittinger K, Chen Y-Y, Keilbaugh SA, Bewtra M, Knights D, Walters WA, Knight R, et al. Linking long-term dietary patterns with gut microbial enterotypes. Science (1979) 2011;334:105–8.

28. Shinn LM, Li Y, Mansharamani A, Auvil LS, Welge ME, Bushell C, Khan NA, Charron CS, Novotny JA, Baer DJ, et al. Fecal bacteria as biomarkers for predicting food intake in healthy adults. J Nutr 2021;151:423–33.

29. Shinn LM, Mansharamani A, Baer DJ, Novotny JA, Charron CS, Khan NA, Zhu R, Holscher HD. Fecal metabolites as biomarkers for predicting food intake by healthy adults. J Nutr 2022;152:2956–65.

30. Novotny JA, Gebauer SK, Baer DJ. Discrepancy between the Atwater factor predicted and empirically measured energy values of almonds in human diets. American Journal of Clinical Nutrition 2012;96:296–301.

31. Edwards CG, Walk AM, Thompson SV, Reeser GE, Erdman JW, Burd NA, Holscher HD, Khan NA. Effects of 12-week avocado consumption on cognitive function among adults with overweight and obesity. Int J Psychophysiol 2020;148:13–24.

32. Thompson SV, Bailey MA, Taylor AM, Kaczmarek JL, Mysonhimer AR, Edwards CG, Reeser GE, Burd NA, Khan NA, Holscher HD. Avocado consumption alters gastrointestinal bacteria abundance and microbial metabolite concentrations among adults with overweight or obesity: A randomized controlled trial. J Nutr 2020;151:753–62.

33. Charron CS, Vinyard BT, Ross SA, Seifried HE, Jeffery EH, Novotny JA. Absorption and metabolism of isothiocyanates formed from broccoli glucosinolates: effects of BMI and daily consumption in a randomised clinical trial. Br J Nutr 2018;120:1370–9.

34. Baer DJ, Gebauer SK, Novotny JA. Walnuts consumed by healthy adults provide less available energy than predicted by the Atwater factors. J Nutr 2016;146:9–13.

35. Thompson SV, Swanson KS, Novotny JA, Baer DJ, Holscher HD. Gastrointestinal microbial changes following whole grain barley and oat consumption in healthy men and women. The FASEB Journal 2016;30:406.1–406.1.

36. Holscher HD, Taylor AM, Swanson KS, Novotny JA, Baer DJ. Almond consumption and processing affects the composition of the gastrointestinal microbiota of healthy adult men and women: A randomized controlled trial. Nutrients 2018;10:126.

37. Kaczmarek JL, Liu X, Charron CS, Novotny JA, Jeffery EH, Seifried HE, Ross SA, Miller MJ, Swanson KS, Holscher HD. Broccoli consumption affects the human gastrointestinal microbiota. Journal of Nutritional Biochemistry 2019;63:27–34.

38. Holscher HD, Guetterman HM, Swanson KS, An R, Matthan NR, Lichtenstein AH, Novotny JA, Baer DJ. Walnut consumption alters the gastrointestinal microbiota, microbially derived secondary bile acids, and health markers in healthy adults: a randomized controlled trial. J Nutr 2018;148:861–7.

39. Rognes T, Flouri T, Nichols B, Quince C, Mahé F. VSEARCH: A versatile open source tool for metagenomics. PeerJ 2016;4:e2584.

40. KneadData – The Huttenhower Lab. Available from: https://huttenhower.sph.harvard.edu/kneaddata/

41. Buchfink B, Xie C, Huson DH. Fast and sensitive protein alignment using DIAMOND. Nat Methods 2014;12:59–60.

42. Sayers EW, Bolton EE, Brister JR, Canese K, Chan J, Comeau DC, Connor R, Funk K, Kelly C, Kim S, et al. Database resources of the national center for biotechnology information. Nucleic Acids Res 2022;50:D20.

43. Huson DH, Beier S, Flade I, Górska A, El-Hadidi M, Mitra S, Ruscheweyh H-J, Tappu R. MEGAN community edition - Interactive exploration and analysis of large-scale microbiome sequencing data. PLoS Comput Biol 2016;12:e1004957.

44. Kanehisa M, Furumichi M, Sato Y, Ishiguro-Watanabe M, Tanabe M. KEGG: integrating viruses and cellular organisms. Nucleic Acids Res 2021;49:D545–51.

45. Kanehisa M. Toward understanding the origin and evolution of cellular organisms. Protein Sci 2019;28:1947–51.

46. Kanehisa M, Goto S. KEGG: Kyoto encyclopedia of genes and genomes. Nucleic Acids Res 2000;28:27–30.

47. Student. The probable error of a mean. Biometrika JSTOR; 1908;6:1.

48. Finotello F, Di Camillo B. Measuring differential gene expression with RNA-seq: challenges and strategies for data analysis. Brief Funct Genomics 2015;14:130–42.

49. Benjamini Y, Hochberg Y. Controlling the false discovery rate: a practical and powerful approach to multiple testing. J R Stat Soc Series B Stat Methodol 1995;57:289–300.

50. limma: linear models for microarray and RNA-seq data. Available from: https://bioinf.wehi.edu.au/limma/

51. Reimand J, Isserlin R, Voisin V, Kucera M, Tannus-Lopes C, Rostamianfar A, Wadi L, Meyer M, Wong J, Xu C, et al. Pathway enrichment analysis and visualization of omics data using g:Profiler, GSEA, Cytoscape and EnrichmentMap. Nat Protoc 2019;14:482–517.

52. Fisher RA. Statistical methods for research workers. Springer, New York, NY; 1992;66–70.

53. Liaw A, Wiener M. Classification and regression by randomForest. R News. 2002. p. 18–22.

54. Buitinck L, Louppe G, Blondel M, Pedregosa F, Müller AC, Grisel O, Niculae V, Prettenhofer P, Gramfort A, Grobler J, et al. API design for machine learning software: Experiences from the scikit-learn project. ECML PKDD 2013.

55. Breiman L, Friedman J, Stone CJ, Olshen RA. Classification and regression trees. Boca Raton, FL: Chapman & Hall/CRC press; 1984.

56. Yilmaz P, Parfrey LW, Yarza P, Gerken J, Pruesse E, Quast C, Schweer T, Peplies J, Ludwig W, Glöckner FO. The SILVA and “all-species living tree project (LTP)” taxonomic frameworks. Nucleic Acids Res 2014;42:643–8.

57. Jackson JE. A user’s guide to principal components. New York: John Wiley & Sons; 2005.

58. McKnight PE, Najab J. Kruskal-Wallis test. The Corsini Encyclopedia of Psychology. 2010. p. 1.

59. Kaoutari AE, Armougom F, Gordon JI, Raoult D, Henrissat B. The abundance and variety of carbohydrate-active enzymes in the human gut microbiota. Nat Rev Microbiol 2013. p. 497– 504.

60. Turnbaugh PJ, Ridaura VK, Faith JJ, Rey FE, Knight R, Gordon JI. The effect of diet on the human gut microbiome: A metagenomic analysis in humanized gnotobiotic mice. Sci Transl Med 2009;1:1–10.

61. Cabezas A, Ribeiro JM, Rodrigues JR, López-Villamizar I, Fernández A, Canales J, Pinto RM, Costas MJ, Cameselle JC. Molecular bases of catalysis and ADP-Ribose preference of human Mn2+-dependent ADP-Ribose/CDP-Alcohol diphosphatase and conversion by mutagenesis to a preferential cyclic ADP-Ribose phosphohydrolase. PLoS One 2015;10:e0118680.

62. Han YH, Garron ML, Kim HY, Kim WS, Zhang Z, Ryu KS, Shaya D, Xiao Z, Cheong C, Kim YS, et al. Structural snapshots of heparin depolymerization by heparin lyase I. J Biol Chem 2009;284:34019.

63. Chimento DP, Mohanty AK, Kadner RJ, Wiener MC. Substrate-induced transmembrane signaling in the cobalamin transporter BtuB. Nat Struct Mol Biol 2003;10:394–401.

64. Galán JE, Lara-Tejero M, Marlovits TC, Wagner S. Bacterial type III secretion systems: specialized nanomachines for protein delivery into target cells. Annu Rev Microbiol 2014;68:415.

65. Walter KA, Bennetti GN, Papoutsakisi ET. Molecular characterization of two Clostridium acetobutylicum ATCC 824 butanol dehydrogenase isozyme genes. J Bacteriol 1992;174:7149–58.

66. David LA, Maurice CF, Carmody RN, Gootenberg DB, Button JE, Wolfe BE, Ling A v, Devlin AS, Varma Y, Fischbach MA, et al. Diet rapidly and reproducibly alters the human gut microbiome. Nature 2014;505:559–63.

67. Harris ZN, Dhungel E, Mosior M, Ahn TH. Massive metagenomic data analysis using abundance-based machine learning. Biol Direct 2019;14:1–13.

68. Yatsunenko T, Rey FE, Manary MJ, Trehan I, Dominguez-Bello MG, Contreras M, Magris M, Hidalgo G, Baldassano RN, Anokhin AP, et al. Human gut microbiome viewed across age and geography. Nature 2012;486:222–7.

69. Raudys SJ, Jain AK. Small sample size effects in statistical pattern recognition: Recommendations for practitioners. IEEE Trans Pattern Anal Mach Intell 1991;13:252–64.

70. Luan J, Zhang C, Xu B, Xue Y, Ren Y. The predictive performances of random forest models with limited sample size and different species traits. Fish Res 2020;227:105534.

71. Morrison DJ, Preston T. Formation of short chain fatty acids by the gut microbiota and their impact on human metabolism. Gut Microbes 2016. p. 189–200.

72. Mukherjee A, Lordan C, Ross RP, Cotter PD. Gut microbes from the phylogenetically diverse genus Eubacterium and their various contributions to gut health. Gut Microbes 2020;12.

73. Liu X, Mao B, Gu J, Wu J, Cui S, Wang G, Zhao J, Zhang H, Chen W. Blautia-a new functional genus with potential probiotic properties? Gut Microbes 2021;13:1–21.

74. Holscher HD, Caporaso JG, Hooda S, Brulc JM, Fahey Jr. GC, Swanson KS. Fiber supplementation influences phylogenetic structure and functional capacity of the human intestinal microbiome: follow-up of a randomized controlled trial. Am J Clin Nutr 2015;101:55–64.

75. Ranjan R, Rani A, Metwally A, McGee HS, Perkins DL. Analysis of the microbiome: Advantages of whole genome shotgun versus 16S amplicon sequencing. Biochem Biophys Res Commun 2016;469:967–77.

76. Pearl J. Causal inference in statistics: An overview. Statist Surv 2009;3:96–146.

77. Imbens GW, Rubin DB. Causal Inference for Statistics, Social, and Biomedical Sciences: An Introduction. Causal Inference: For Statistics, Social, and Biomedical Sciences an Introduction. New York: Cambridge University Press; 2015.

78. Huttenhower C, Gevers D, Knight R, Abubucker S, Badger JH, Chinwalla AT, Creasy HH, Earl AM, Fitzgerald MG, Fulton RS, et al. Structure, function and diversity of the healthy human microbiome. Nature 2012;486:207–14.

79. Lee BY, Ordovás JM, Parks EJ, Anderson CAM, Barabási A-L, Clinton SK, de la Haye K, Duffy VB, Franks PW, Ginexi EM, et al. Research gaps and opportunities in precision nutrition: an NIH workshop report. Am J Clin Nutr 2022;116(6):1877–1900.

