## Supplemental Material for "Fecal metagenomics to identify biomarkers of food intake in healthy adults: Findings from randomized, controlled, nutrition trials"

**Online Supporting Material (OSM)**

**Supplemental Table 1. Nutrient compositions of control and treatment provisions for healthy adult participants across avocado, almond, broccoli, and walnut trials[1].**

| **Nutrient** | **Avocado**(31)[2] | |
|---|---|---|
| | **Control** | **Avocado** |
| **Energy (kcal/day)** | | |
| **Males** | 660 | 662 |
| **Females** | 538 | 530 |
| **Total fat (% of energy)** | 39 | 40 |
| *Saturated* | 17 | 7 |
| *Monounsaturated* | 10 | 24 |
| *Polyunsaturated* | 9 | 5 |
| **Carbohydrate (% of energy)** | 45 | 45 |
| **Total dietary fiber (g/day)** | 4 | 16 |
| *Soluble fiber* | 1 | 6 |
| *Pectins* | 0 | 4 |
| *Insoluble fiber* | 3 | 10 |
| **Protein (% of energy)** | 16 | 14 |

| **Nutrient** | **Almond**(30) | | **Broccoli**(33) | | **Walnut**(34) | |
|---|---|---|---|---|---|---|
| | **Control** | **Almond** | **Control** | **Broccoli** | **Control** | **Walnut** |
| **Energy (kcal/day)** | Assigned based on Harris Benedict equation | | Met individual energy requirements for weight maintenance | | Met individual energy requirements for weight maintenance | |
| **Fat (% of energy)** | 25 | 30 | 30 | Identical to control + 200 g or broccoli and 20 g of daikon radish/day* | 29 | Reduced in equal proportions during the walnut treatment condition to achieve isocaloric food intake† |
| **Carbohydrate (% of energy)** | 59 | 55 | 54 | | 54 | |
| **Dietary fiber from specific intervention food (%DV)** | - | 15 | - | 19 | - | 10 |
| **Protein (% of energy)** | 16 | 15 | 16 | * | 17 | † |

[1] Whole grains nutrient compositions have been excluded due to unpublished data
[2] Values provided are for one provided, controlled meal/day (not total intake levels)

**Supplemental Table 2. Top 50 differentially abundant KEGG orthologs in almond, broccoli, and walnut treatment groups compared to their respective controls.** For each food group, differential abundance analysis was conducted by contrasting the normalized KO counts of each treatment group against its corresponding control group using Student's t-test. The Benjamini-Hochberg procedure was used to control for the false discovery rate. KOs were considered differentially abundant if they met a significance threshold of $q < 0.20$.

| Almond Treatment vs. Control | | Broccoli Treatment vs. Control | | Walnut Treatment vs. Control | |
|---|---|---|---|---|---|
| **KO** | **q** | **KO** | **q** | **KO** | **q** |
| K11912 | 0.018 | K02843 | 0.0000 | K10562 | 0.006 |
| K18023 | 0.018 | K13695 | 0.0000 | K16013 | 0.010 |
| K03634 | 0.038 | K21344 | 0.0001 | K10560 | 0.010 |
| K03851 | 0.038 | K04762 | 0.0001 | K10559 | 0.010 |
| K00437 | 0.038 | K10121 | 0.0001 | K11494 | 0.010 |
| K01501 | 0.052 | K08223 | 0.0001 | K14188 | 0.010 |
| K13683 | 0.055 | K02858 | 0.0001 | K00100 | 0.015 |
| K00856 | 0.055 | K03718 | 0.0001 | K02122 | 0.016 |
| K11742 | 0.061 | K02066 | 0.0001 | K03779 | 0.017 |
| K07151 | 0.065 | K09011 | 0.0001 | K03780 | 0.017 |
| K06951 | 0.065 | K15778 | 0.0001 | K13573 | 0.017 |
| K08172 | 0.065 | K01960 | 0.0001 | K02574 | 0.020 |
| K11384 | 0.092 | K03113 | 0.0001 | K11031 | 0.021 |
| K11473 | 0.092 | K00946 | 0.0001 | K20490 | 0.021 |
| K21947 | 0.116 | K00342 | 0.0001 | K10118 | 0.021 |
| K00573 | 0.122 | K01798 | 0.0001 | K01420 | 0.021 |
| K13017 | 0.122 | K02065 | 0.0001 | K10615 | 0.025 |
| K12950 | 0.122 | K09882 | 0.0001 | K04758 | 0.025 |
| K13821 | 0.130 | K06076 | 0.0001 | K05881 | 0.025 |
| K09857 | 0.130 | K01843 | 0.0001 | K10561 | 0.026 |
| K01517 | 0.134 | K00347 | 0.0001 | K05588 | 0.026 |
| K19303 | 0.148 | K13598 | 0.0001 | K21600 | 0.026 |
| K10107 | 0.148 | K21345 | 0.0001 | K00198 | 0.026 |
| K03224 | 0.148 | K01719 | 0.0001 | K20491 | 0.026 |
| K13695 | 0.154 | K12510 | 0.0001 | K08234 | 0.026 |
| K18008 | 0.163 | K03270 | 0.0001 | K23536 | 0.026 |
| K03219 | 0.163 | K03775 | 0.0001 | K03205 | 0.026 |
| K11743 | 0.163 | K03694 | 0.0001 | K00334 | 0.026 |
| K06151 | 0.168 | K00174 | 0.0001 | K00887 | 0.026 |
| K19714 | 0.168 | K00175 | 0.0001 | K14138 | 0.026 |
| K06192 | 0.174 | K06020 | 0.0001 | K11050 | 0.028 |
| K23301 | 0.174 | K18138 | 0.0001 | K22015 | 0.029 |
| K19050 | 0.174 | K01144 | 0.0001 | K19050 | 0.029 |
| K18011 | 0.174 | K02230 | 0.0001 | K19092 | 0.029 |
| K19200 | 0.174 | K00992 | 0.0001 | K04028 | 0.029 |
| K01561 | 0.174 | K18302 | 0.0001 | K09779 | 0.029 |
| K03226 | 0.174 | K03089 | 0.0001 | K07657 | 0.029 |
| K12146 | 0.174 | K17204 | 0.0001 | K09251 | 0.030 |
| K16011 | 0.183 | K18954 | 0.0001 | K03486 | 0.033 |
| K05805 | 0.183 | K01840 | 0.0001 | K19077 | 0.033 |
| K01093 | 0.183 | K16089 | 0.0001 | K10119 | 0.033 |
| K07492 | 0.183 | K03562 | 0.0001 | K07738 | 0.033 |
| K09688 | 0.183 | K06861 | 0.0001 | K20488 | 0.033 |
| K11957 | 0.183 | K00350 | 0.0001 | K03706 | 0.033 |
| K03148 | 0.183 | K09808 | 0.0001 | K02435 | 0.033 |
| K12542 | 0.183 | K01082 | 0.0001 | K15023 | 0.033 |
| K08305 | 0.183 | K00176 | 0.0001 | K18350 | 0.033 |
| K01295 | 0.183 | K01308 | 0.0001 | K18348 | 0.035 |
| K03567 | 0.183 | K11621 | 0.0001 | K04027 | 0.035 |
| K07019 | 0.183 | K06142 | 0.0001 | K09145 | 0.035 |

**Supplemental Table 3. Differentially abundant pathways in almond treatment vs. control.** Pathway analysis was performed on the set of differentially abundant KOs ($q < 0.20$) using the *limma* R package. Pathways with an uncorrected $P < 0.05$ were considered differentially abundant. This table describes the ID, name, number of total KOs, number of differentially abundant KOs, and *P* value of the significance of each pathway.

| **Pathway ID** | **Pathway Name** | **N** | **DE** | **P.DE** |
|---|---|---:|---:|---:|
| path:ko03070 | Bacterial secretion system | 72 | 4 | 9.34E-05 |
| path:ko00541 | O-Antigen nucleotide sugar biosynthesis | 70 | 2 | 2.34E-02 |
| path:ko00311 | Penicillin and cephalosporin biosynthesis | 9 | 1 | 3.03E-02 |
| path:ko02025 | Biofilm formation - Pseudomonas aeruginosa | 84 | 2 | 3.29E-02 |

**Supplemental Table 4. Differentially abundant pathways in broccoli treatment vs. control.** Pathway analysis was performed on the set of differentially abundant KOs ($q < 0.20$) using the *limma* R package. Pathways with an uncorrected $P < 0.05$ were considered differentially abundant. This table describes the ID, name, number of total KOs, number of differentially abundant KOs, and $P$ value of the significance of each pathway.

| Pathway ID | Pathway | N | DE | P.DE |
|---|---|---|---|---|
| path:ko01100 | Metabolic pathways | 3288 | 781 | 8.07E-35 |
| path:ko02010 | ABC transporters | 494 | 167 | 1.38E-19 |
| path:ko02020 | Two-component system | 455 | 140 | 8.65E-13 |
| path:ko02060 | Phosphotransferase system (PTS) | 71 | 39 | 1.02E-12 |
| path:ko01230 | Biosynthesis of amino acids | 231 | 85 | 1.22E-12 |
| path:ko00520 | Amino sugar and nucleotide sugar metabolism | 146 | 58 | 1.60E-10 |
| path:ko01110 | Biosynthesis of secondary metabolites | 1107 | 264 | 6.00E-09 |
| path:ko01240 | Biosynthesis of cofactors | 352 | 104 | 1.09E-08 |
| path:ko01250 | Biosynthesis of nucleotide sugars | 151 | 55 | 1.90E-08 |
| path:ko00720 | Carbon fixation pathways in prokaryotes | 90 | 38 | 3.43E-08 |
| path:ko00051 | Fructose and mannose metabolism | 108 | 43 | 3.53E-08 |
| path:ko00620 | Pyruvate metabolism | 123 | 47 | 3.76E-08 |
| path:ko01200 | Carbon metabolism | 315 | 93 | 7.03E-08 |
| path:ko00540 | Lipopolysaccharide biosynthesis | 58 | 28 | 7.04E-08 |
| path:ko01120 | Microbial metabolism in diverse environments | 911 | 218 | 1.34E-07 |
| path:ko00550 | Peptidoglycan biosynthesis | 53 | 25 | 6.20E-07 |
| path:ko00650 | Butanoate metabolism | 98 | 37 | 1.49E-06 |
| path:ko00010 | Glycolysis / Gluconeogenesis | 99 | 37 | 1.98E-06 |
| path:ko00500 | Starch and sucrose metabolism | 97 | 36 | 3.28E-06 |
| path:ko00640 | Propanoate metabolism | 91 | 34 | 5.16E-06 |
| path:ko00541 | O-Antigen nucleotide sugar biosynthesis | 70 | 28 | 7.71E-06 |
| path:ko00020 | Citrate cycle (TCA cycle) | 57 | 24 | 1.22E-05 |
| path:ko03070 | Bacterial secretion system | 72 | 27 | 4.40E-05 |
| path:ko00270 | Cysteine and methionine metabolism | 106 | 35 | 8.00E-05 |
| path:ko00130 | Ubiquinone and other terpenoid-quinone biosynthesis | 52 | 21 | 8.86E-05 |
| path:ko00531 | Glycosaminoglycan degradation | 14 | 9 | 1.34E-04 |
| path:ko00400 | Phenylalanine, tyrosine and tryptophan biosynthesis | 66 | 24 | 2.01E-04 |
| path:ko01232 | Nucleotide metabolism | 108 | 34 | 2.84E-04 |
| path:ko00121 | Secondary bile acid biosynthesis | 10 | 7 | 3.73E-04 |
| path:ko00511 | Other glycan degradation | 20 | 10 | 9.09E-04 |
| path:ko02024 | Quorum sensing | 211 | 55 | 1.13E-03 |
| path:ko00920 | Sulfur metabolism | 90 | 28 | 1.16E-03 |
| path:ko00552 | Teichoic acid biosynthesis | 24 | 11 | 1.26E-03 |
| path:ko01502 | Vancomycin resistance | 21 | 10 | 1.46E-03 |
| path:ko00052 | Galactose metabolism | 67 | 22 | 1.76E-03 |
| path:ko00030 | Pentose phosphate pathway | 77 | 24 | 2.47E-03 |
| path:ko00230 | Purine metabolism | 172 | 45 | 2.84E-03 |
| path:ko01503 | Cationic antimicrobial peptide (CAMP) resistance | 51 | 17 | 4.76E-03 |
| path:ko00290 | Valine, leucine and isoleucine biosynthesis | 17 | 8 | 4.83E-03 |
| path:ko00240 | Pyrimidine metabolism | 96 | 27 | 6.75E-03 |
| path:ko00450 | Selenocompound metabolism | 29 | 11 | 7.47E-03 |
| path:ko00190 | Oxidative phosphorylation | 183 | 45 | 9.59E-03 |
| path:ko00340 | Histidine metabolism | 42 | 14 | 9.98E-03 |
| path:ko00730 | Thiamine metabolism | 34 | 12 | 1.01E-02 |
| path:ko00680 | Methane metabolism | 165 | 41 | 1.09E-02 |
| path:ko00900 | Terpenoid backbone biosynthesis | 47 | 15 | 1.20E-02 |
| path:ko00630 | Glyoxylate and dicarboxylate metabolism | 93 | 25 | 1.61E-02 |
| path:ko04122 | Sulfur relay system | 28 | 10 | 1.68E-02 |
| path:ko00750 | Vitamin B6 metabolism | 17 | 7 | 1.93E-02 |
| path:ko00220 | Arginine biosynthesis | 54 | 16 | 2.01E-02 |
| path:ko00250 | Alanine, aspartate and glutamate metabolism | 64 | 18 | 2.42E-02 |
| path:ko00571 | Lipoarabinomannan (LAM) biosynthesis | 14 | 6 | 2.43E-02 |
| path:ko00330 | Arginine and proline metabolism | 87 | 23 | 2.46E-02 |
| path:ko00521 | Streptomycin biosynthesis | 15 | 6 | 3.47E-02 |
| path:ko01210 | 2-Oxocarboxylic acid metabolism | 63 | 17 | 4.08E-02 |
| path:ko00260 | Glycine, serine and threonine metabolism | 96 | 24 | 4.12E-02 |
| path:ko00572 | Arabinogalactan biosynthesis - Mycobacterium | 12 | 5 | 4.45E-02 |
| path:ko02026 | Biofilm formation - Escherichia coli | 50 | 14 | 4.52E-02 |
| path:ko00780 | Biotin metabolism | 20 | 7 | 4.77E-02 |

**Supplemental Table 5. Differentially abundant pathways in walnut treatment vs. control.** Pathway analysis was performed on the set of differentially abundant KOs ($q < 0.20$) using the *limma* R package. Pathways with an uncorrected $P < 0.05$ were considered differentially abundant. This table describes the ID, name, number of total KOs, number of differentially abundant KOs, and p value of the significance of each pathway.

| Pathway ID | Pathway | N | DE | P.DE |
|---|---|---|---|---|
| path:ko02010 | ABC transporters | 494 | 79 | 1.19E-20 |
| path:ko02060 | Phosphotransferase system (PTS) | 71 | 24 | 3.72E-14 |
| path:ko01502 | Vancomycin resistance | 21 | 9 | 4.05E-07 |
| path:ko00051 | Fructose and mannose metabolism | 108 | 20 | 5.00E-07 |
| path:ko02024 | Quorum sensing | 211 | 26 | 2.98E-05 |
| path:ko00052 | Galactose metabolism | 67 | 13 | 3.04E-05 |
| path:ko01120 | Microbial metabolism in diverse environments | 911 | 73 | 6.46E-05 |
| path:ko02020 | Two-component system | 455 | 43 | 7.31E-05 |
| path:ko00520 | Amino sugar and nucleotide sugar metabolism | 146 | 19 | 1.72E-04 |
| path:ko00620 | Pyruvate metabolism | 123 | 17 | 1.81E-04 |
| path:ko01100 | Metabolic pathways | 3288 | 201 | 7.35E-04 |
| path:ko00552 | Teichoic acid biosynthesis | 24 | 6 | 1.11E-03 |
| path:ko00550 | Peptidoglycan biosynthesis | 53 | 9 | 1.40E-03 |
| path:ko00010 | Glycolysis / Gluconeogenesis | 99 | 13 | 1.65E-03 |
| path:ko01200 | Carbon metabolism | 315 | 29 | 1.69E-03 |
| path:ko00720 | Carbon fixation pathways in prokaryotes | 90 | 12 | 2.16E-03 |
| path:ko00470 | D-Amino acid metabolism | 47 | 7 | 9.79E-03 |
| path:ko01054 | Nonribosomal peptide structures | 29 | 5 | 1.53E-02 |
| path:ko00920 | Sulfur metabolism | 90 | 10 | 1.73E-02 |
| path:ko01501 | beta-Lactam resistance | 66 | 8 | 1.97E-02 |
| path:ko00500 | Starch and sucrose metabolism | 97 | 10 | 2.76E-02 |
| path:ko00730 | Thiamine metabolism | 34 | 5 | 2.90E-02 |
| path:ko00640 | Propanoate metabolism | 91 | 9 | 4.47E-02 |
| path:ko01503 | Cationic antimicrobial peptide (CAMP) resistance | 51 | 6 | 4.64E-02 |

**Supplemental Table 6. Top 10 KEGG orthologs (KOs) with feature importance scores (FIS) extracted from the random forest models classifying almond treatment vs. control, broccoli treatment vs. control, and walnut treatment vs. control.** Random forest models trained in a leave-one-out fashion were used to examine the relationship between food consumption and change in KO abundance. Only KOs that were differentially abundant (*q* < 0.20) were included in the model.

| **Almond Treatment vs. Control** | |
|---|---|
| **KO** | **FIS** |
| K12950 ctpC; manganese-transporting P-type ATPase C [EC:7.2.2.22] | 0.075 |
| K13683 wcaE; putative colanic acid biosynthesis glycosyltransferase [EC:2.4.-.-] | 0.060 |
| K01501 E3.5.5.1; nitrilase [EC:3.5.5.1] | 0.051 |
| K18023 mbhJ; membrane-bound hydrogenase subunit mbhJ [EC:1.12.7.2] | 0.048 |
| K03089 rpoH; RNA polymerase sigma-32 factor | 0.044 |
| K11473 glcF; glycolate oxidase iron-sulfur subunit | 0.040 |
| K13695 nlpC; probable lipoprotein NlpC | 0.032 |
| K11912 ppkA; serine/threonine-protein kinase PpkA [EC:2.7.11.1] | 0.032 |
| K13821 putA; RHH-type transcriptional regulator, proline utilization regulon repressor / proline dehydrogenase / delta 1-pyrroline-5-carboxylate dehydrogenase [EC:1.5.5.2 1.2.1.88] | 0.030 |
| K03634 lolA; outer membrane lipoprotein carrier protein | 0.030 |
| **Broccoli Treatment vs. Control** | |
| K02843 waaF, rfaF; heptosyltransferase II [EC:2.4.-.-] | 0.023 |
| K13695 nlpC; probable lipoprotein NlpC | 0.020 |
| K01093 appA; 4-phytase / acid phosphatase [EC:3.1.3.26 3.1.3.2] | 0.014 |
| K08223 fsr; MFS transporter, FSR family, fosmidomycin resistance protein | 0.013 |
| K01423 bepA; beta-barrel assembly-enhancing protease [EC:3.4.-.-] | 0.012 |
| K03270 kdsC; 3-deoxy-D-manno-octulosonate 8-phosphate phosphatase (KDO 8-P phosphatase) [EC:3.1.3.45] | 0.011 |
| K11537 xapB; MFS transporter, NHS family, xanthosine permease | 0.011 |
| K21344 rfaE1; D-glycero-beta-D-manno-heptose-7-phosphate kinase [EC:2.7.1.167] | 0.011 |
| K16692 etk-wzc; tyrosine-protein kinase Etk/Wzc [EC:2.7.10.3] | 0.010 |
| K15778 pmm-pgm; phosphomannomutase / phosphoglucomutase [EC:5.4.2.8 5.4.2.2] | 0.010 |
| **Walnut Treatment vs. Control** | |
| K10562 rhaT; rhamnose transport system ATP-binding protein [EC:7.5.2.-] | 0.041 |
| K16013 cydD; ATP-binding cassette, subfamily C, bacterial CydD | 0.040 |
| K10559 rhaS; rhamnose transport system substrate-binding protein | 0.038 |
| K06336 tasA, cotN; spore coat-associated protein N | 0.031 |
| K09145 K09145; uncharacterized protein | 0.026 |
| K03779 ttdA; L(+)-tartrate dehydratase alpha subunit [EC:4.2.1.32] | 0.025 |
| K11031 slo; thiol-activated cytolysin | 0.024 |
| K19050 hepA; heparin lyase [EC:4.2.2.7] | 0.023 |
| K05658 ABCB1, CD243; ATP-binding cassette, subfamily B (MDR/TAP), member 1 [EC:7.6.2.2] | 0.023 |
| K11494 RCBTB; RCC1 and BTB domain-containing protein | 0.022 |

**Supplemental Table 7. Top 25 KEGG orthologs (KOs) with feature importance scores (FIS) extracted from the random forest almond treatment vs. broccoli treatment vs. walnut treatment multi-food model.** Random forest models trained in a leave-one-out fashion were used to examine the relationship between food consumption and change in KO abundance. Only KOs that were differentially abundant ($q < 0.20$) were included in the model.

| Almond Treatment vs. Broccoli Treatment vs. Walnut Treatment | |
|---|---|
| **KO** | **FIS** |
| K10974 codB; cytosine permease | 0.0086 |
| K13683 wcaE; putative colanic acid biosynthesis glycosyltransferase [EC:2.4.-.-] | 0.0081 |
| K08676 tri; tricorn protease [EC:3.4.21.-] | 0.0079 |
| K00335 nuoF; NADH-quinone oxidoreductase subunit F [EC:7.1.1.2] | 0.0076 |
| K00965 galT, GALT; UDPglucose--hexose-1-phosphate uridylyltransferase [EC:2.7.7.12] | 0.0070 |
| K03270 kdsC; 3-deoxy-D-manno-octulosonate 8-phosphate phosphatase (KDO 8-P phosphatase) [EC:3.1.3.45] | 0.0064 |
| K12510 tadB; tight adherence protein B | 0.0063 |
| K02533 lasT; tRNA/rRNA methyltransferase [EC:2.1.1.-] | 0.0061 |
| K19050 hepA; heparin lyase [EC:4.2.2.7] | 0.0059 |
| K08305 mltB; membrane-bound lytic murein transglycosylase B [EC:4.2.2.-] | 0.0057 |
| K11957 natA; neutral amino acid transport system ATP-binding protein | 0.0057 |
| K04028 eutN; ethanolamine utilization protein EutN | 0.0056 |
| K00176 korD, oorD; 2-oxoglutarate ferredoxin oxidoreductase subunit delta [EC:1.2.7.3] | 0.0054 |
| K11954 natB; neutral amino acid transport system substrate-binding protein | 0.0053 |
| K04056 yscO, sctO; type III secretion protein O | 0.0051 |
| K03779 ttdA; L(+)-tartrate dehydratase alpha subunit [EC:4.2.1.32] | 0.0051 |
| K18023 mbhJ; membrane-bound hydrogenase subunit mbhJ [EC:1.12.7.2] | 0.0049 |
| K05838 ybbN; putative thioredoxin | 0.0049 |
| K00334 nuoE; NADH-quinone oxidoreductase subunit E [EC:7.1.1.2] | 0.0046 |
| K14260 alaA; alanine-synthesizing transaminase [EC:2.6.1.66 2.6.1.2] | 0.0046 |
| K09797 K09797; uncharacterized protein | 0.0046 |
| K04027 eutM; ethanolamine utilization protein EutM | 0.0045 |
| K00437 hydB; [NiFe] hydrogenase large subunit [EC:1.12.2.1] | 0.0045 |
| K01241 amn; AMP nucleosidase [EC:3.2.2.4] | 0.0044 |
| K10559 rhaS; rhamnose transport system substrate-binding protein | 0.0044 |

**Supplemental Table 8. Prediction of specific food intake from adults who consumed 5 foods using random forest using NCBI-nr metagenomic data and SILVA 16S annotated taxonomic data.**

| | Almond, n | Avocado, n | Broccoli, n | Walnut, n | Whole Grains, n | Balanced Accuracy, % |
|---|---|---|---|---|---|---|
| Almond | | | | | | |
| Metagenomic | 6 | 2 | 5 | 0 | 2 | 69 |
| 16S | 4 | 2 | 2 | 4 | 3 | 62 |
| Metagenomic control | 0 | 9 | 2 | 1 | 3 | 44 |
| 16S control | 0 | 10 | 3 | 0 | 2 | 49 |
| Avocado | | | | | | |
| Metagenomic | 2 | 19 | 0 | 5 | 0 | 80 |
| 16S | 3 | 15 | 4 | 1 | 3 | 74 |
| Metagenomic control | 3 | 16 | 0 | 1 | 0 | 71 |
| 16S control | 1 | 14 | 2 | 2 | 1 | 68 |
| Broccoli | | | | | | |
| Metagenomic | 0 | 0 | 12 | 0 | 3 | 87 |
| 16S | 0 | 5 | 9 | 0 | 1 | 77 |
| Metagenomic control | 1 | 12 | 0 | 2 | 0 | 48 |
| 16S control | 0 | 9 | 3 | 1 | 2 | 56 |
| Walnut | | | | | | |
| Metagenomic | 0 | 5 | 0 | 13 | 0 | 83 |
| 16S | 0 | 1 | 0 | 13 | 4 | 83 |
| Metagenomic control | 2 | 1 | 1 | 7 | 7 | 66 |
| 16S control | 0 | 3 | 0 | 2 | 13 | 52 |
| Whole Grains | | | | | | |
| Metagenomic | 0 | 3 | 0 | 0 | 27 | 92 |
| 16S | 0 | 0 | 0 | 1 | 29 | 91 |
| Metagenomic control | 2 | 2 | 0 | 0 | 11 | 79 |
| 16S control | 0 | 0 | 0 | 1 | 14 | 83 |
| Overall Accuracy and AUC | | | | | | |
| Metagenomic | | | | | | 74 (AUC: 0.93) |
| 16S | | | | | | 67 (AUC: 0.89) |
| Metagenomic control | | | | | | 40 (AUC: 0.68) |
| 16S control | | | | | | 40 (AUC: 0.67) |

**Supplemental Table 9. Microbial biomarkers using the top fecal 16S SILVA annotated microbiota species from metabolically healthy adults who consumed 5 foods.**

| Rank | Overall variable importance | SILVA assignment |
|---|---|---|
| 1 | 0.034 | *Roseburia* undefined |
| 2 | 0.026 | *Roseburia* spp. |
| 3 | 0.015 | Undefined species in Rikenellaceae family |
| 4 | 0.011 | *Roseburia* spp. |
| 5 | 0.010 | *Ruminococcus torques* group undefined |
| 6 | 0.009 | *Ruminiclostridium* spp. |
| 7 | 0.009 | *Bacteroides* spp. |
| 8 | 0.009 | *Parabacteroides* spp. |
| 9 | 0.009 | *Alistipes* spp. |
| 10 | 0.008 | Family XIII *UCG-001* undefined |
| 11 | 0.007 | Family XIII *UCG-001* spp. |
| 12 | 0.007 | *Subdoligranulum* spp. |
| 13 | 0.006 | *Subdoligranulum* spp. |
| 14 | 0.006 | Ruminococcaceae *UCG-003* undefined |
| 15 | 0.005 | *Veillonellaceae* spp. |
| 16 | 0.005 | Undefined species in Family XIII family |
| 17 | 0.004 | Lachnospiraceae *UCG-001* undefined |
| 18 | 0.003 | Ruminococcaceae *UCG-010* undefined |
| 19 | 0.003 | Undefined species in Firmicutes phylum |
| 20 | 0.003 | *Sutterella* spp. |
| 21 | 0.003 | Undefined species in Clostridiales vadin BB60 group family |
| 22 | 0.003 | *Sutterella* spp. |
| 23 | 0.002 | Ruminococcaceae *UCG-009* undefined |
| 24 | 0.002 | Ruminococcaceae *UCG-013* undefined |
| 25 | 0.002 | Undefined species in Lachnospiraceae |
